# Copper as a central element critical to interkingdom interactions within a polymicrobial wound environment

**DOI:** 10.64898/2026.02.10.705071

**Authors:** Samuel Fenn, Mario Recker, Ruth C. Massey

**Affiliations:** School of Microbiology, University College Cork, Cork, Ireland; APC Microbiome Ireland, University College Cork, Cork, Ireland; Centre for Ecology and Conservation, University of Exeter, Penryn Campus, United Kingdom; Institute for Tropical Medicine, University of Tübingen, Tübingen, Germany; School of Cellular and Molecular Medicine, University of Bristol, Bristol, UK

## Abstract

*Staphylococcus aureus* is a global health concern, estimated to cause over one million deaths each year. To colonise and survive in humans, *S. aureus* must overcome the host-immune system and compete with other native and invading microorganisms. As a frequent coloniser of wounds, it forms inter-kingdom polymicrobial biofilms, which are associated with impaired wound healing and poorer clinical outcomes. However, the microbial processes that contribute to the competitiveness of *S. aureus* within infection-related inter-kingdom communities are poorly understood. To address this, we established a polymicrobial *in vitro* wound-like model (WLM) consisting of two common bacterial pathogens *Staphylococcus aureus* and *Pseudomonas aeruginosa,* as well as a the fungal pathobiont *Candida albicans*. Using a collection of clinical *S. aureus* strains we then applied a genome-wide association study (GWAS) approach to identify bacterial factors that contribute to *S. aureus* fitness within this environment. Amongst the six *S. aureus* loci identified was *copA*, which encodes a copper exporter that prevents the accumulation of copper in the bacterial cytoplasm. Characterising the molecular processes involved, we identified a 4-way interaction, where *C. albicans* releases copper from the ceruloplasmin found in the human host’s plasma, which in turn causes a build-up of copper within the *S. aureus* cytoplasm, rendering the bacteria more susceptible to killing by *P. aeruginosa*. Human infection data validates the clinical relevance of this, where there is an association between mutations in *copA* and an origin of infection that is wound-based, highlighting the importance of understanding the entire infection ecosystem if improvements therapeutic approaches are to be developed.

## INTRODUCTION

*Staphylococcus aureus* is a significant global health concern with an estimated 1.1 million deaths attributed to this pathogen annually (1). This opportunistic pathogen can cause a myriad of diseases at diverse sites in the host including wound, respiratory, urinary, bone and bloodstream infections (2). However, the most common association of *S. aureus* with humans is as part of the microbiota operating as a commensal in the anterior nares, with up to a third of people colonised with few ill effects (3,4). Whether acting as a commensal or a pathogen, these lifestyles and the varying infection sites represent drastically different challenges to *S. aureus,* and its continuing prevalence highlights how its adaptability contributes to its global success.

Wound infections represent a major healthcare burden with management of this disease type estimated to cost the USA $148.65 billion dollars in 2022 alone (5). The wound microenvironment is polymicrobial in nature, colonised by normal skin commensals such as *S. aureus* and other opportunistic pathogens including *Pseudomonas aeruginosa*, *Escherichia coli*, *Klebsiella pneumoniae*, *Enterococcus* sp., and the dimorphic fungus *Candida albicans* (6–9). In previous work we have applied genome-wide association studies (GWAS) to characterise various aspects of *S. aureus* infection biology (10–16), where phenotype were quantified with *S. aureus* grown as mono-species cultures. However, to establish itself at different host infection sites this pathogen must outcompete the resident microbiota and other invading organisms, and the processes governing it success in these polymicrobial environments are poorly understood. Existence within such environments has been shown to alter the behaviours of pathogens (17–21), where wounds coinfected with *S. aureus* & *C. albicans* or *S. aureus* & *P. aeruginosa* are associated with increased healing times and poorer clinical outcomes (22). In both instances, poorer clinical outcomes are associated with increased biofilm formation, resulting in higher levels of antimicrobial tolerance when compared to mono-species infections (18,20–24).

To address the knowledge gap surrounding colonisation and competition of *S. aureus* in polymicrobial communities, we adapted a previously characterised *in vitro* wound-like model (WLM), which allows co-culture of *S. aureus*, *C. albicans*, and *P. aeruginosa* in multi-species biofilms (25). Using this model, we determined the fitness of 134 clinical *S. aureus* isolates when in competition with *C. albicans* and *P. aeruginosa*. A GWAS identified six novel effectors, which impact *S. aureus* colonisation and competition in poly-microbial communities, including the copper exporter CopA. We further characterised the role of CopA, and demonstrate that copper acquired from the host can be used by one competing pathogen to make *S. aureus* susceptible to killing by another competing pathogen, and that the ability to control intracellular copper levels is critical to the success of *S. aureus* in polymicrobial wound environments.

## RESULTS

### Wound-like model establishment and validation

To identify factors that contribute to the ability of *S. aureus* to successfully compete within a poly-microbial wound environment we established an *in vitro* Wound-Like Model (WLM), that has been shown previously by Deleon and colleagues (2014) to have utility when investigating interactions between *S. aureus* and *P. aeruginosa* (25). To determine the effect of *C. albicans* addition to the previously established *S. aureus* & *P. aeruginosa* model, mono-species and polymicrobial cultures were set up and the density of each microorganism determined by measuring colony forming units per millilitre of the medium (CFU/mL) over seven days. A visible biofilm was evident, when *S. aureus* was present within the community (Fig. 1A). Mono-species cultures in both WLM and Tryptic Soy Broth (TSB) achieved stable populations across the seven days with *S. aureus* reaching 10^9^, *P. aeruginosa* 10^10^ and *C. albicans* 10^7^ CFU/mL (Fig 1B). In a polymicrobial WLM environment a 1-log reduction in CFU/mL recovery was exhibited by all organisms at early stages, followed by a further 1-log reduction in CFU/mL across the 7-day monitoring period when compared to mono-species cultures (Fig 1C). Despite the reduction, a stable and reproducible polymicrobial community was established with 10^7^ *S. aureus*, 10^8^ *P. aeruginosa* and 10^5^ *C. albicans* CFU/mL after seven days.

**Figure 1.**
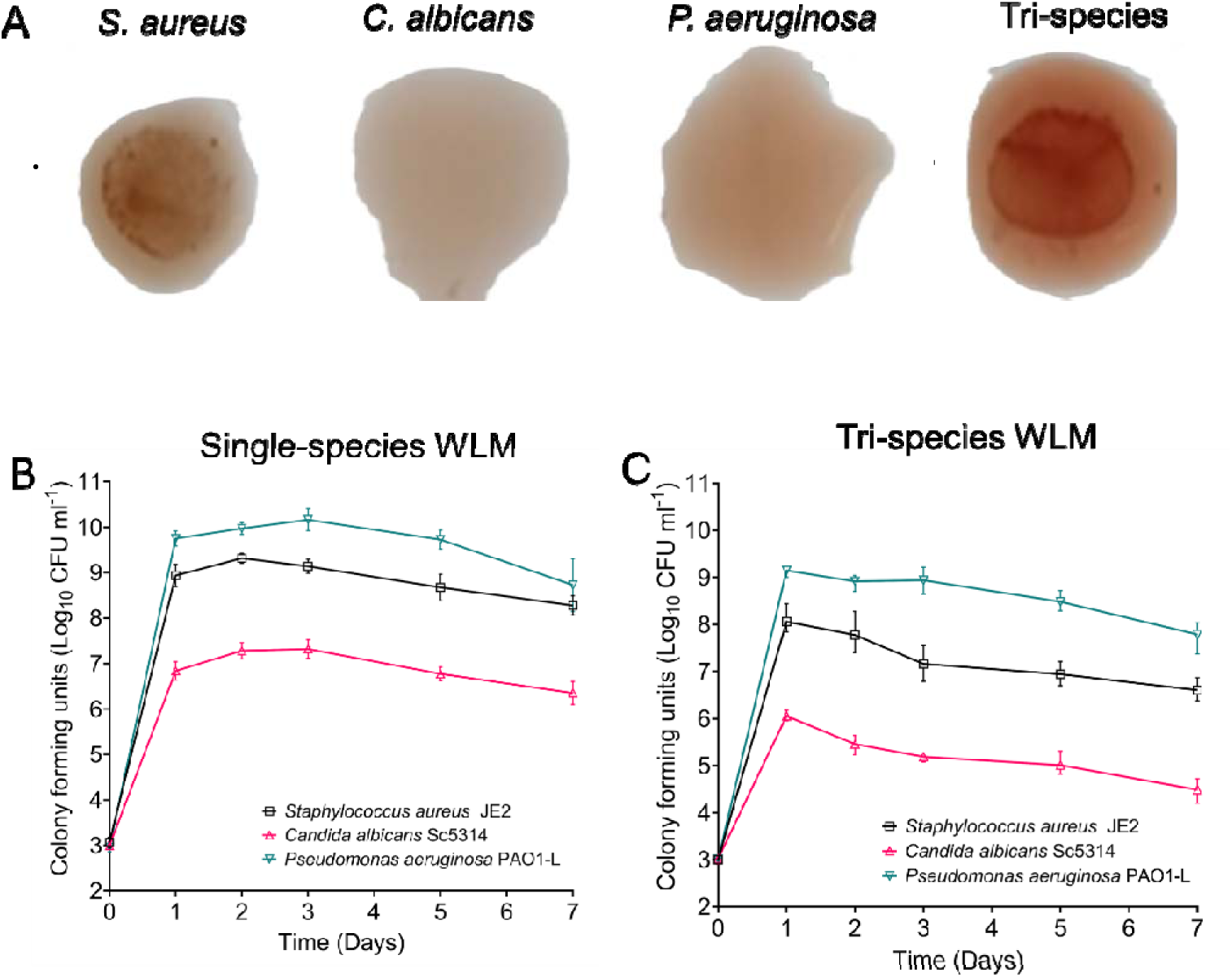
Establishment and validation of an interkingdom wound-like model containing *S. aureus* JE2, *P. aeruginosa* PAO1-L and *C. albicans* Sc5314. (**A**) Photograph of WLM biofilms at seven days, with visible biofilm formation only observed when *S. aureus* is present (indicated by arrows). (**B**) Time-course of community composition in mono-species and (**C**) polymicrobial cultures through CFU recovery on selective media. Data presented is the mean of three independent experiments with error bars representing the standard deviation.

### Clinical *S. aureus* strains demonstrate varying abilities to compete in polymicrobial wound-like biofilms

A collection of 134 clinical *S. aureus* isolates were used to assess fitness within the polymicrobial WLM. Each clinical isolate was cultured in this model for three days, and the density of *S. aureus* enumerated (CFU/mL) as a measure of relative fitness (Fig 2a). Significant variability was found across the collection with a 2-log variance in density between the least and the most fit *S. aureus* isolates (Fig 2a). The genome sequence for all 134 isolates were available, which allowed us to identify novel loci or SNPs (single nucleotide polymorphisms) contributing to fitness within the poly-microbial WLM by means of a genome wide association study (GWAS). This identified 42 loci that were significantly associated with altered fitness within the WLM (Table S1). Of these, 32 were available for functional validation from the Nebraska transposon mutant library (Table S1) (26), which we used to establish both mono and polymicrobial biofilms in the WLM. Eight mutants demonstrated significantly altered fitness in the polymicrobial model (Fig S1), whilst no impact was observed in mono-species cultures (Fig S1), suggesting that their impact is solely on *S. aureus* competitive fitness within the polymicrobial WLM.

**Figure 2.**
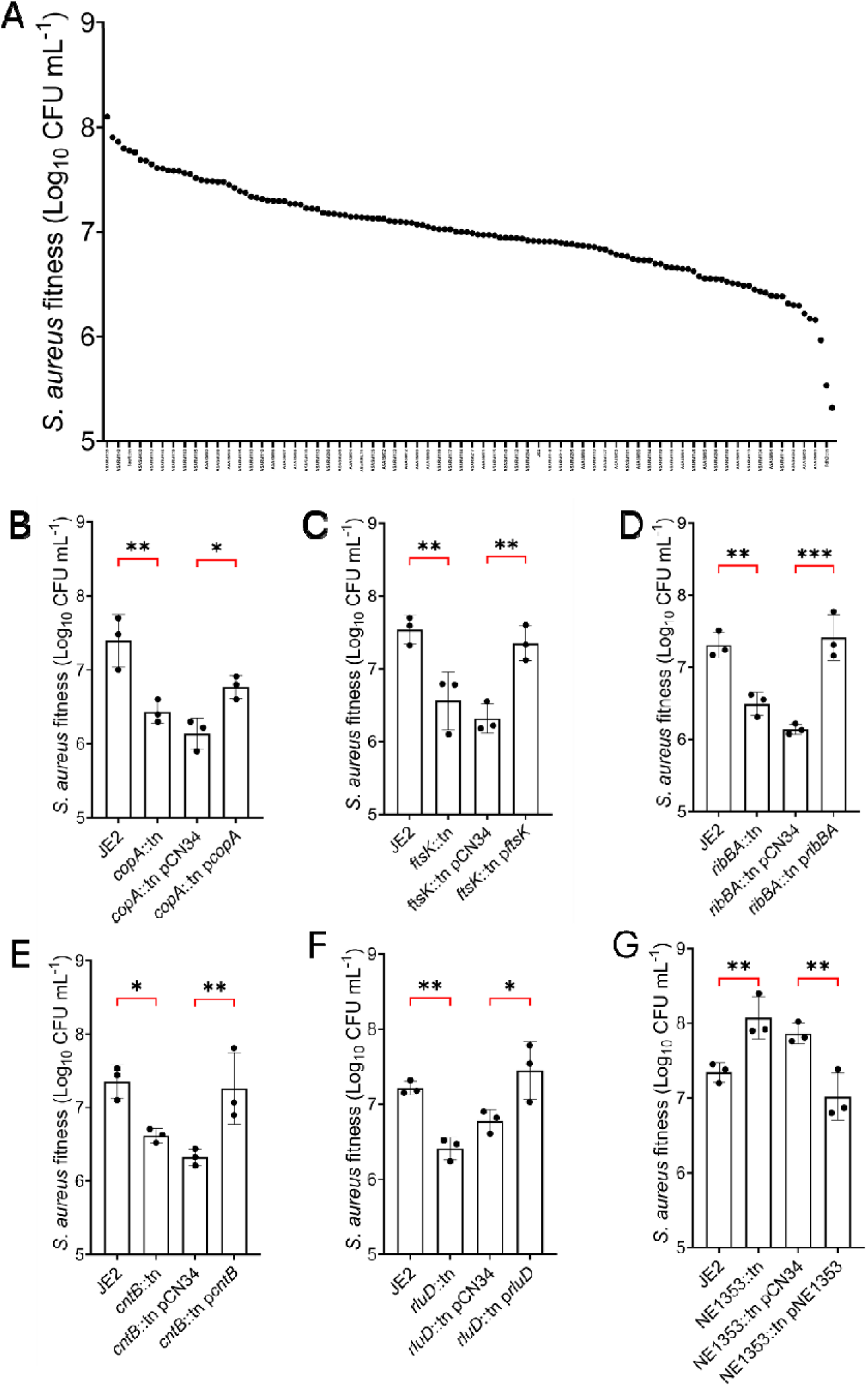
*S. aureus* fitness in the polymicrobial WLM is variable and poly-genic in nature. (A) Density as a measure of relative fitness of 134 CC22 clinical *S. aureus* isolates when co-cultured alongside *P. aeruginosa* and *C. albicans* in the WLM (CFU/mL). (B-G) Functional validation of the GWAS identified fitness associated loci. Of the original 42 identified loci, six were validated as affecting the phenotype by both mutation and complementation of the fitness phenotype: *copA* (B), *ftsK* (C), *ribBA* (D), *cntB* (E), *rluD* (F), and NE1353 (G). Recovery of *P. aeruginosa* and *C. albicans* can be found in supplementary (Table S2). Each dot represents an individual experiment, the bars represent the mean value, error bars the standard deviation. Significance determined as *<0.05, **<0.01, ***<0.001 calculated by one-way ANOVA with Sidak’s (B-G) multiple comparisons test.

We then used the pCN34 plasmid to complement each mutation by expressing the gene *in trans,* which restored the fitness of six of the eight identified mutants (Fig 2B-FG). The six genes were the copper ATPase transporter *copA* (Fig 2C); DNA translocase *ftsK* (Fig 2D); riboflavin biosynthesis protein *ribBA* (Fig 2E); staphylopine mediated metal ion transporter *opp1B*/*cntB* (Fig 2F); pseudouridine ribosomal RNA modification gene *rluD* (Fig 2G); and an amidase-domain containing protein SAUSA300_2579 (Fig 2H). The two genes we were unable to complement were the sex pheromone *camS* and an amidohydrolase encoding gene SAUSA300_0534.

### A single polymorphism in *copA* is associated with improved fitness in multi-species WLM

Copper is both a beneficial micronutrient and a potent antimicrobial at higher concentrations, inducing bacterial killing through multiple redox mediated mechanisms such as ROS generation and protein/enzyme damage (27,28). Balancing copper homeostasis is therefore critical, with the CopA protein playing a central role in this as an ATP-dependant transporter responsible for removing excess copper from the bacterial cytoplasm (29,30). Whilst the role of CopA has previously been explored in *S. aureus* monocultures (29,30), its role in interkingdom interactions has not yet been established. The polymorphism in *copA* identified by our GWAS approach was a 1G>A mutation replacing the atypical start codon GTG with the more commonly used ATG. Ten of the isolates in the collection contained this allele of *copA,* which was associated with an increase in fitness in the polymicrobial WLM (Fig 3A). We hypothesised that this substitution may result in enhanced translation of CopA. To test this we constructed a translational reporter using *gfp* fused to a DNA fragment containing the promoter and the first 10 amino acids of CopA, under the control of either the GTG or the ATG start codon. Like many exporter systems, the expression of this gene is induced by the presence of its substrate, and in accordance with this, minimal GFP induction was observed in the absence of copper (Fig 3B). However, when cultured in the presence of 200 μM copper both GTG- and ATG-driven production of GFP was observed, with expression from the ATG codon demonstrating higher fluorescence (Fig 3B), indicative of higher translational activity.

**Figure 3.**
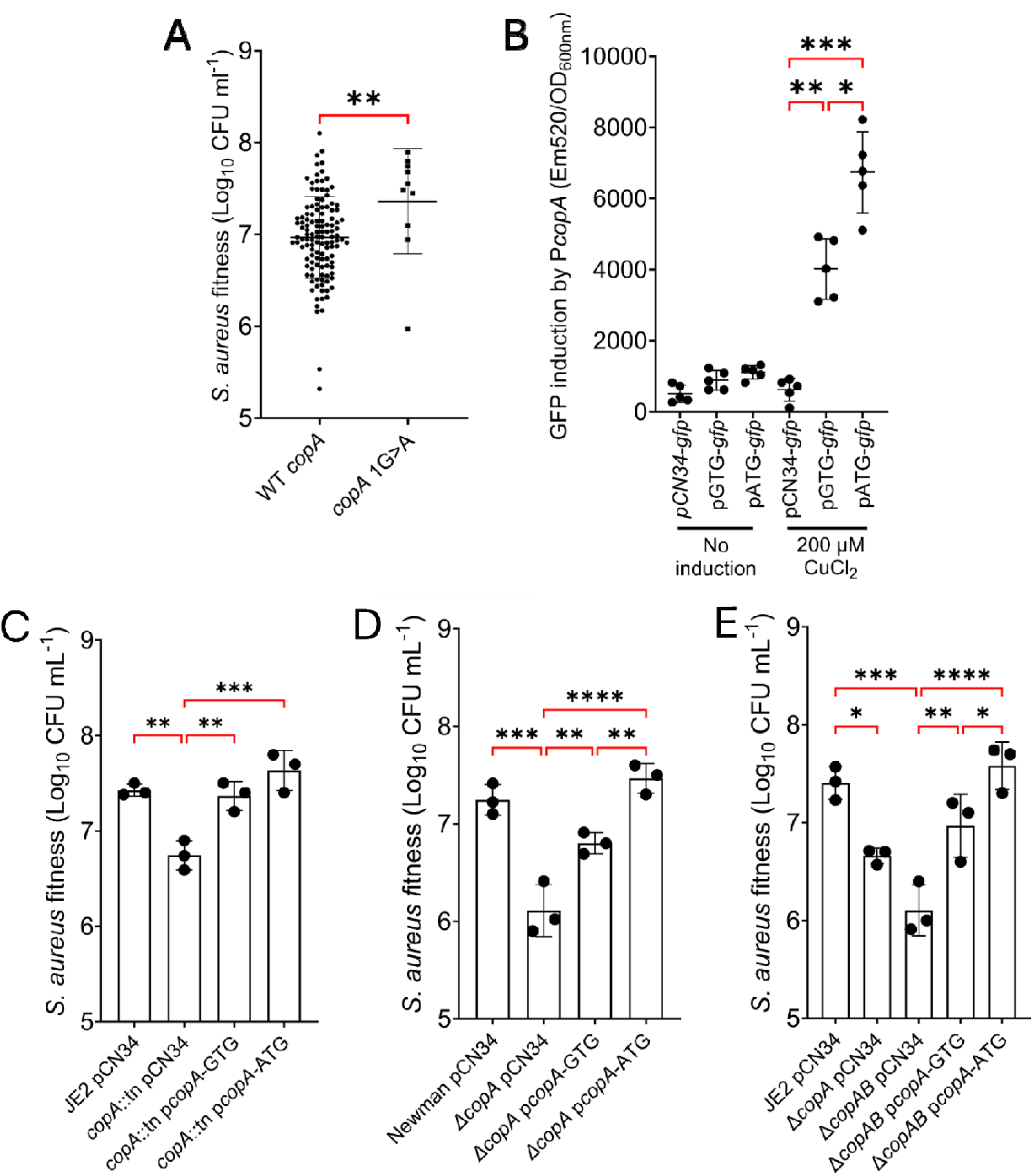
Impact of *copA* mutant allele on *S. aureus* fitness in a polymicrobial WLM. (A) Fitness of the *copA* SNP containing clinical isolates relative to those without the SNP. (B-D). Fitness of *S. aureus* strains at day 3 in the polymicrobial WLM. (B) In the JE2 copA mutant background, (C) is in the Newman *copA* mutant background and (D) is in the JE2 *copAB* double mutant background. Bars represent the mean value of three biological replicates, error bars the standard deviation. Significance determined as *<0.05, **<0.01, ***<0.001, ****<0.0001 calculated using an unpaired t-test (A), and Tukey’s (B) or Sidak’s (D and E) multiple comparisons test.

To confirm whether higher translational activity equates to increased protein activity, we generated a second *copA* complementation plasmid, this time with *copA* translation under the control of the alternate ATG start codon (p*copA*-ATG), and compared its phenotype-complementing ability to that of the wild type GTG start codon containing plasmid (p*copA*-GTG). Initially, these plasmids were transformed into JE2 *copA*::tn, but they both complemented the fitness phenotype to equivalent levels (Fig 3C). However, community acquired MRSA strains, such as JE2, possess a secondary copper resistance operon *copBL,* with *copB* producing a secondary copper transporter (29), that may mask any effect of the change in start codon of *copA*. To account for this, a deletion mutant of *copA* was constructed in *S. aureus* strain Newman, which only has the *copA* copper transporter system. We also mutated both *copA* and *copB* in the JE2 background and transduced our two complementing plasmids into both mutants. The fitness complementing ability of both p*copA*-GTG and p*copA*-ATG was then compared in the polymicrobial WLM. Newman Δ*copA* demonstrating a larger decrease in fitness when compared to JE2 Δ*copA.* (Fig 3D-E). Mutation of both *copA* and *copB* in the JE2 background further reduced fitness in the WLM relative to the *copA* mutant alone. Complementation of the Newman *copA* mutant and JE2 double *copAB* mutant with p*copA*-ATG demonstrated enhanced fitness in comparison to p*copA*-GTG in the polymicrobial WLM (Fig 3D-E), verifying the beneficial effect the ATG variant has over the GTG expressing one in this environment.

To understand the dynamics of the fitness effects, we monitored *S. aureus copA* mutants over seven days in both JE2 and Newman backgrounds. No significant difference in fitness was found at day 1 when compared to the wild type strain. However, a reduction in fitness for the copper exporter mutants was observed in subsequent days as the WLM matured (Fig S2). This indicates that the loss in competitive fitness is not due to a growth defect, as they have reached peak density at this point, but rather an inability to compete and survive in this polymicrobial community. Furthermore, equivalent *S. aureus*-only WLM biofilms do not demonstrate this effect, indicating the observed phenotype is due to the polymicrobial nature of this model (Fig S2).

### The reduced fitness of the *copA* mutant is dependent on both *C. albicans* and *P. aeruginosa*

To determine the interspecies interaction responsible for the altered fitness of *S. aureus* in the polymicrobial WLM, we measured its density over subsequent days as mono, dual and polymicrobial models. In the mono-species WLM, mutation of *copA* did not impact the fitness of *S. aureus* (Fig 4A), whereas the loss of *copA* had a small but non-significant effect during dual culture with *P. aeruginosa* (Fig 4B). When cultured as a dual-species biofilm with *C. albicans,* mutation of *copA* resulted in a small, statistically significant reduction in fitness, with complementation restoring this defect (Fig 4C). However, it is in the polymicrobial model in the presence of both *C. albicans* and *P. aeruginosa* where the *copA* mutant is severely compromised, with a 1.4-log reduction in density at day 3 compared to the isogenic wild-type (Fig 4D). Complementation with p*copA-*ATG allele increased fitness of *S. aureus* when compared to p*copA*-GTG in this model (Fig 4D). A similar impact was seen when replicating these experiments using *S. aureus* JE2, with the loss of both copper exporters reducing competitiveness in a polymicrobial community (Fig S2). The relatively small effect exhibited in dual-species biofilms indicates that neither *C. albicans* nor *P. aeruginosa* are solely responsible for the reduced fitness exhibited by *S. aureus* copper exporter mutants. Instead, it suggests that these two organisms act synergistically to compromise the competitiveness of *S. aureus* in the polymicrobial environment.

**Figure 4.**
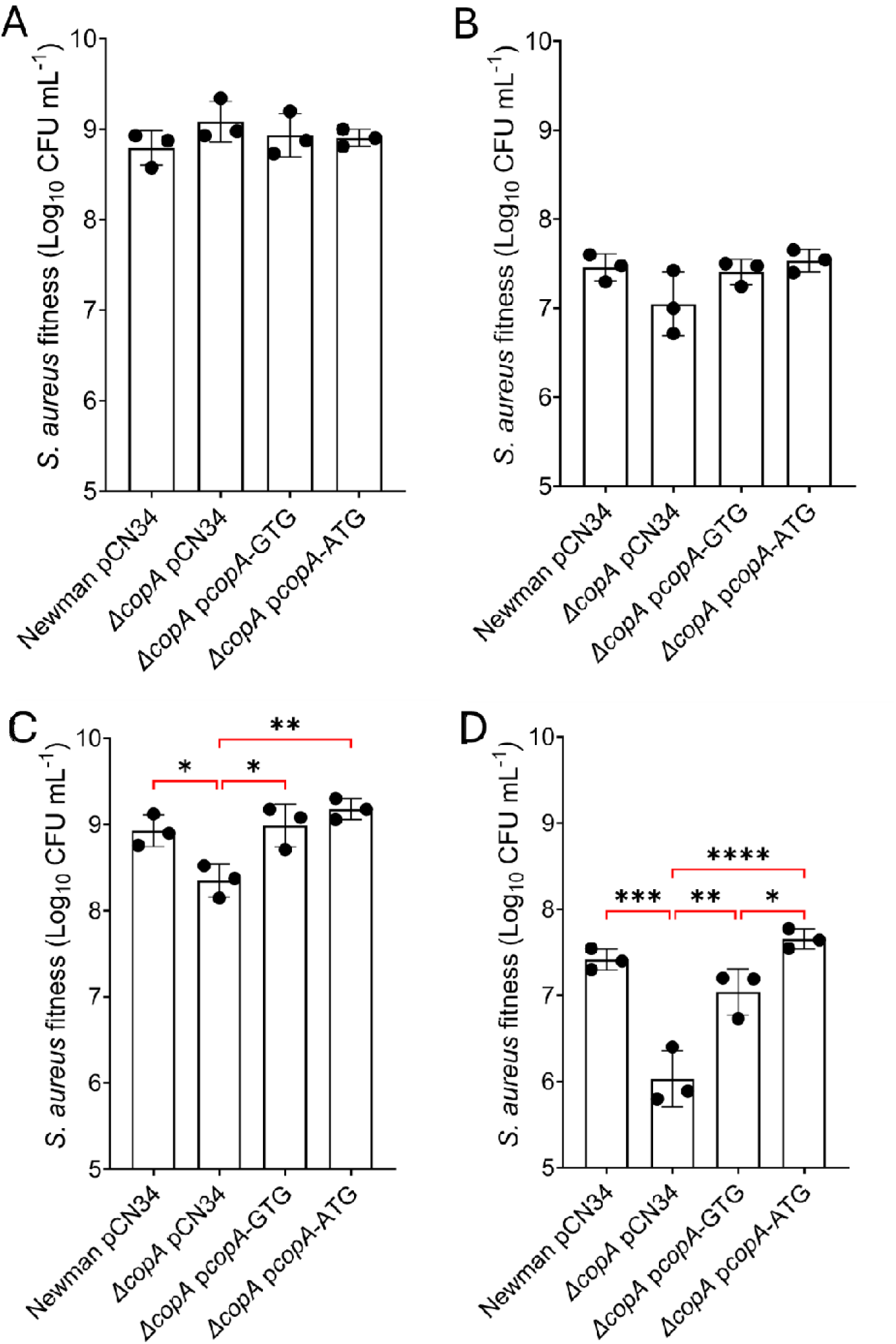
Both *P. aeruginosa* and *C. albicans* are required for the full loss of fitness of the *S. aureus copA* mutant in the polymicrobial WLM. WLM biofilms established with (A) *S. aureus*, (B) *S. aureus* & *P. aeruginosa*, (C) *S. aureus* & *C. albicans*, and (D) *S. aureus* & *P. aeruginosa* & *C. albicans*. Bars represent the mean value of three biological replicates, error bars the standard deviation. Significance determined as *<0.05, **<0.01, ***<0.001, ****<0.0001 calculated using Sidak’s multiple comparisons test.

### *C. albicans* dependent copper release drives accumulation in *S. aureus* copper exporter mutants

Given the defined role of CopA as a copper exporter we hypothesized that copper accumulation in the *S. aureus* cytoplasm is driving the reduced competitiveness of the *copA* mutant in polymicrobial communities. To verify this, mono, dual and polymicrobial WLM biofilms were washed with PBS and treated with lysostaphin to release the intracellular contents of *S. aureus* whilst leaving *P. aeruginosa* and *C. albicans* intact. The copper content of these lysates were then quantified. In *S. aureus* only and *S. aureus* & *P. aeruginosa* WLM biofilms there was no significant increase in copper accumulation in the Newman *copA* mutant (Fig 5A). However, when *S. aureus* was cultured alongside *C. albicans* in WLM, the *copA* mutant accumulated 4-fold more copper in both dual and polymicrobial cultures (Fig 5A). This same pattern was exhibited by JE2 copper exporter mutants with double mutation of *copAB* displaying elevated concentrations of copper in a *C. albicans* dependant manner (Fig S3).

**Figure 5.**
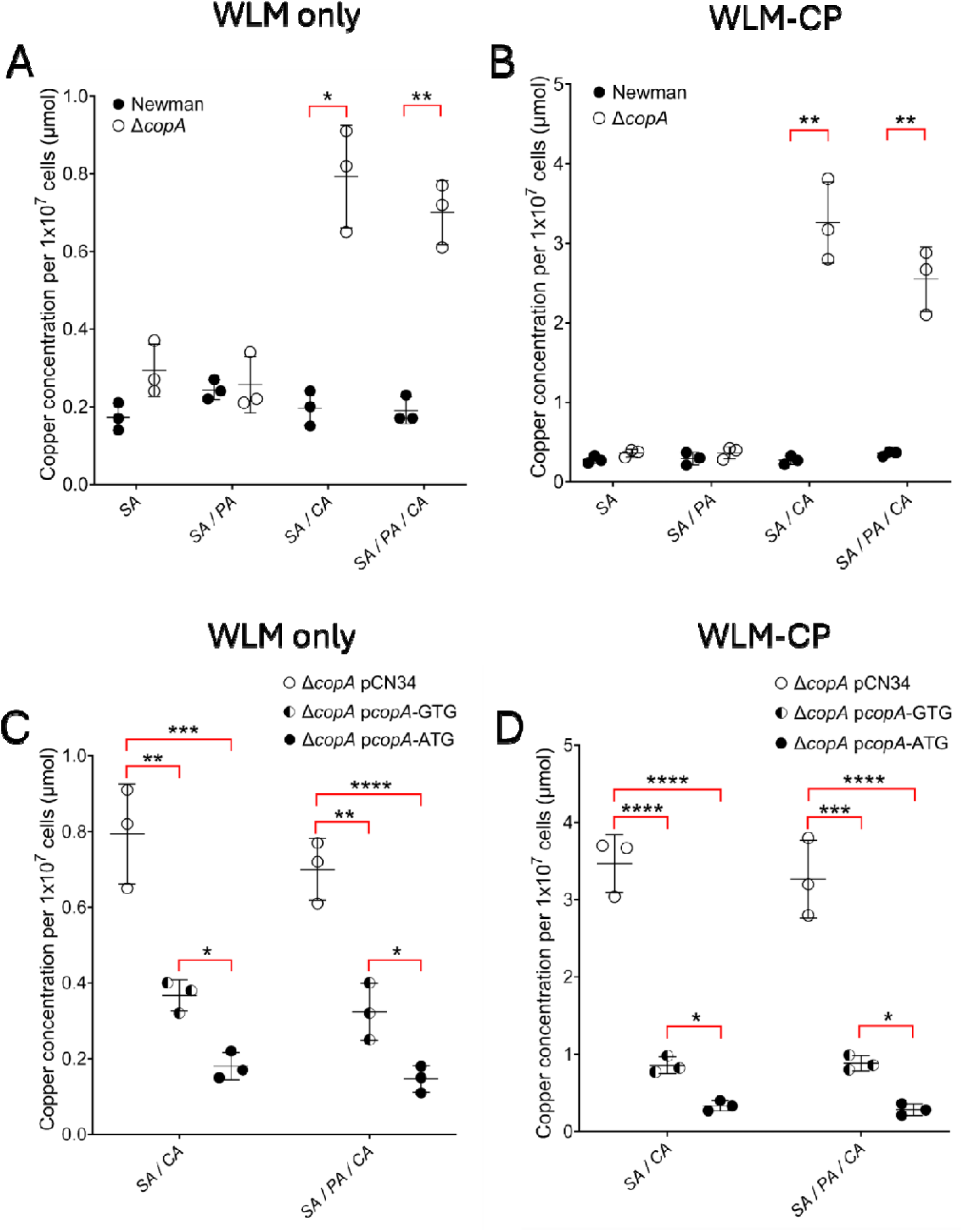
Copper liberated from ceruloplasmin by *C. albicans* accumulates in the *copA* mutants and reduces the fitness of *S. aureus*. (**A**) The copper content of the *S. aureus* cells in the WLM established as mono, dual and tri-species cultures in WLM only and (**B**) where ceruloplasmin has been added (WLM-CP). (**C**) Impact of complementation with the wild type *copA* allele and SNP in WLM or (**D**) in WLM-CP. Lines represent the mean of three biological replicates, error bars the standard deviation. Significance determined as *<0.05, **<0.01, ***<0.001, ****<0.0001 calculated using Sidak’s (A-D) and Tukey’s (E) multiple comparisons test.

Copper is elevated at infections sites through the accumulation of the human copper-binding protein ceruloplasmin (31–33). Besold and colleagues (2021) previously showed that *C. albicans* can release and then utilise copper from ceruloplasmin in serum (34), whilst *S. aureus* and *P. aeruginosa* are not reported to have this activity. The WLM contains 50% plasma, with an additional 5% whole blood, both serving as a source of ceruloplasmin from which *C. albicans* can liberate copper. As it is not possible to remove the ceruloplasmin from the plasma or blood, we determined its effect on the fitness of the *S. aureus copA* mutant by supplementing the WLM with additional exogenous human ceruloplasmin and measuring intracellular copper concentration in mono, dual and tri-species WLM cultures (WLM-CP). In agreement with our previous experiments, copper accumulation in the *copA* mutant is *C. albicans*-dependent (Figure 5B); however, the addition of ceruloplasmin enhanced this effect, with Newman Δ*copA* demonstrating a 10-fold increase in intracellular copper content when established as a dual or tri-species model. Complementation of Newman Δ*copA* with the p*copA*-GTG partially restored intracellular copper concentrations in both WLM and WLM-CP (Figure 5C-D). This same pattern was exhibited by the JE2 *copAB* double mutant, resulting in increased copper accumulation in a *C. albicans* and ceruloplasmin dependent manner (Figure S3A-B). Complementation with p*copA*-ATG resulted in further reductions in cytoplasmic copper levels in both Newman and JE2 backgrounds (Figure 5C-D and S3C-D). Copper toxicity could therefore partially explain the reduced fitness of the *S. aureus copA* mutant in the presence of *C. albicans*, but does not explain the enhanced effect in the additional presence of *P. aeruginosa* (Figure 4C-D).

### Copper accumulation sensitises *S. aureus* to *P. aeruginosa* produced redox active antimicrobials

Copper accumulation increases the susceptibility of *S. aureus* to reactive oxygen species and redox active molecules, with copper acting as a catalyst in Fenton-like chemistry (35). This results in increased production of hydroxyl and superoxide radicals, damaging cells through oxidation of proteins, lipids, fatty acids, and nucleic acids (27,28,36). Given the effect that *C. albicans* has on increasing intracellular copper levels of *S. aureus*, and that *P. aeruginosa* uses redox active antimicrobials to compete in multi-species environments (38–40), we hypothesized that these factors in combination affect *S. aureus* fitness.

To test this we first verified that the increased intracellular copper in the *copA* mutants sensitised *S. aureus* to oxidative stress through exposure to hydrogen peroxide (Fig S4). *P. aeruginosa* produces pyocyanin, hydrogen cyanide (CN) and 2-Heptyl-4-hydroxyquinoline N-oxide (HQNO), with these antimicrobials shown to inhibit *S. aureus* and promote production of ROS through inhibition of the aerobic respiratory chain (38–40). Both pyocyanin and cyanide also inhibit the enzymes catalase and super-oxide dismutase (41–44), mechanisms employed by *S. aureus* to detoxify H_2_O_2_ and O_2_ ^-^ respectively. This led us to hypothesize that *P. aeruginosa* produced antimicrobials that amplify endogenous ROS production and inhibit detoxification mechanisms of *S. aureus*, enhance the toxic effect of *C. albicans* derived copper accumulation exhibited in this model.

To determine if amplification of endogenous ROS is responsible for the loss of *S. aureus* viability we performed pyocyanin, KCN and HQNO sensitivity assays. Polymicrobial WLMs were established with and without ceruloplasmin, and after three days the biofilms were disrupted and normalised to 1×10^6^ *S. aureus* cells in TSB. Strains were then exposed to 0.2 mM pyocyanin, 1 mM KCN, or 0.2 mM HQNO and plated to determine *S. aureus* survival. The Newman *copA* mutant in the WLM showed increased sensitivity to pyocyanin and KCN (Fig 6A-B). The addition of ceruloplasmin and subsequent copper accumulation further increased this sensitivity, demonstrating that loss of copper exporter function sensitises *S. aureus* to *Pseudomonas* produced antimicrobials. Complementation with p*copA*-ATG enhanced resistance to both pyocyanin and KCN when compared to p*copA*-GTG (Figure 6A-B), confirming that increased copper accumulation correlates with enhanced sensitivity to these redox active antimicrobials. Similarly established mono-species biofilms containing only *S. aureus* demonstrated no defect in pyocyanin, KCN killing and HQNO survival (Fig S5). Pre-treatment of polymicrobial biofilms cultured in the presence of WLM-CP with the ROS scavenger thiourea inhibited pyocyanin and KCN mediated killing (Fig 6C-D). This demonstrates that increased ROS generation through the combined action of *C. albicans* induced copper accumulation and *P. aeruginosa* produced pyocyanin and KCN is responsible for the reduced fitness phenotype of *S. aureus* copper exporter mutants in this multi-species biofilm model. Mutation of *copA* also enhanced activity of HQNO; however, the impact of ceruloplasmin was minor and complementation with p*copA*-ATG demonstrated no competitive advantage (Fig S4). Thiourea treatment did not restore this killing phenotype, indicating that the HQNO mediated killing in a *copA* mutant is not ROS mediated (Fig S4).

**Figure 6.**
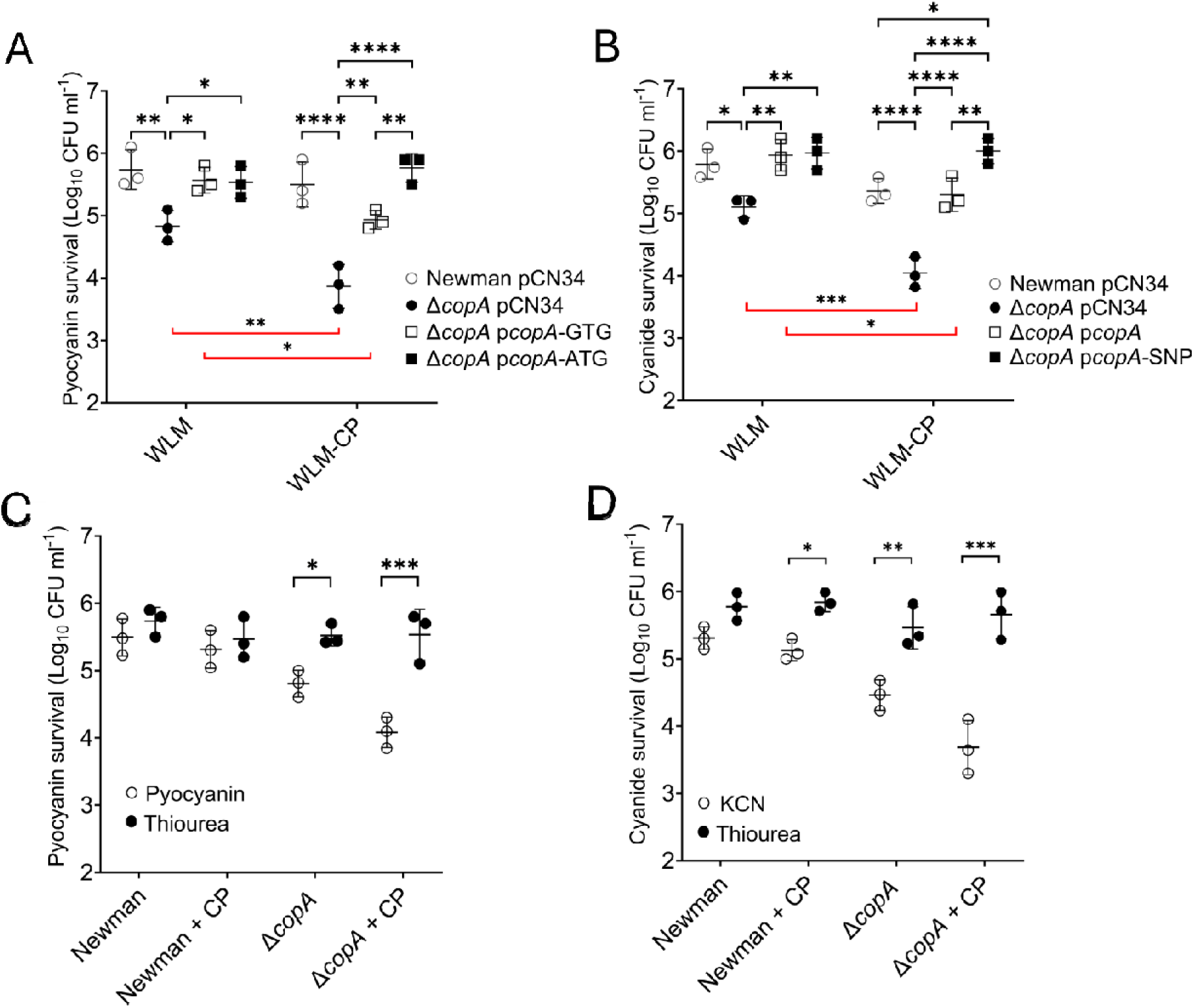
Copper accumulation drives increased sensitivity to *P. aeruginosa* produced antimicrobials. (A) Pyocyanin killing assay of *S. aureus* strains cultured in WLM and WLM-CP as a polymicrobial biofilm. (B) KCN killing assay of *S. aureus* strains cultured in WL and WLM-CP as a polymicrobial biofilm. (C & D) Treatment with the ROS scavenger thiourea protects *S. aureus* from pyocyanin and KCN mediated killing. Bars represent the mean of three independent experiments, error bars the standard deviation. Significance determined as *<0.05, **<0.01, ***<0.001, ****<0.0001 calculated using a two-way Anova with Tukey’s post-hoc test (A-B) and Three-way Anova with Sidak’s multiple comparisons test (C-D)

### CopA alleles affect fitness in environmentally specific ways and have significant clinical associations

The existence of both *copA* alleles within a population of clinical isolates suggests that there are natural environments where they each confer an advantage to the bacteria, preventing the ATG allele from reaching fixation. Up to this point, our measure of fitness has been based on competitive outcomes following co-culture with *P. aeruginosa* and *C. albicans*. However, to understand the prevalence of the GTG allele, we needed to directly measure the within-species competitiveness of both alleles. To do this we established an assay where the fitness of strains was measured relative to a marked rifampicin resistant *S. aureus* strain, which uses the GTG *copA* allele. The wild type Newman, the *copA* mutant, and the p*copA*-ATG or p*copA*-GTG complemented strains were each competed against the marked *S. aureus* in the polymicrobial WLM with and without additional ceruloplasmin. The fitness of the *copA* mutant was significantly impaired in both environments, where the addition of ceruloplasmin further decreased its fitness (Fig 7a). This effect was complemented by expressing either *copA* allele, with p*copA*-ATG increasing competitive fitness when compared to p*copA*-GTG (Fig 7a). This trend was further enhanced in the presence of supplemented ceruloplasmin at concentrations relevant to infection levels (Fig 7b). No competitive advantage was observed in *S. aureus* only WLM monocultures (Fig S6). Together, these findings verify the fitness benefit of the ATG allele in WLM, with increased copper availability through ceruloplasmin supplementation increasing the fitness benefit of this allele.

**Figure 7.**
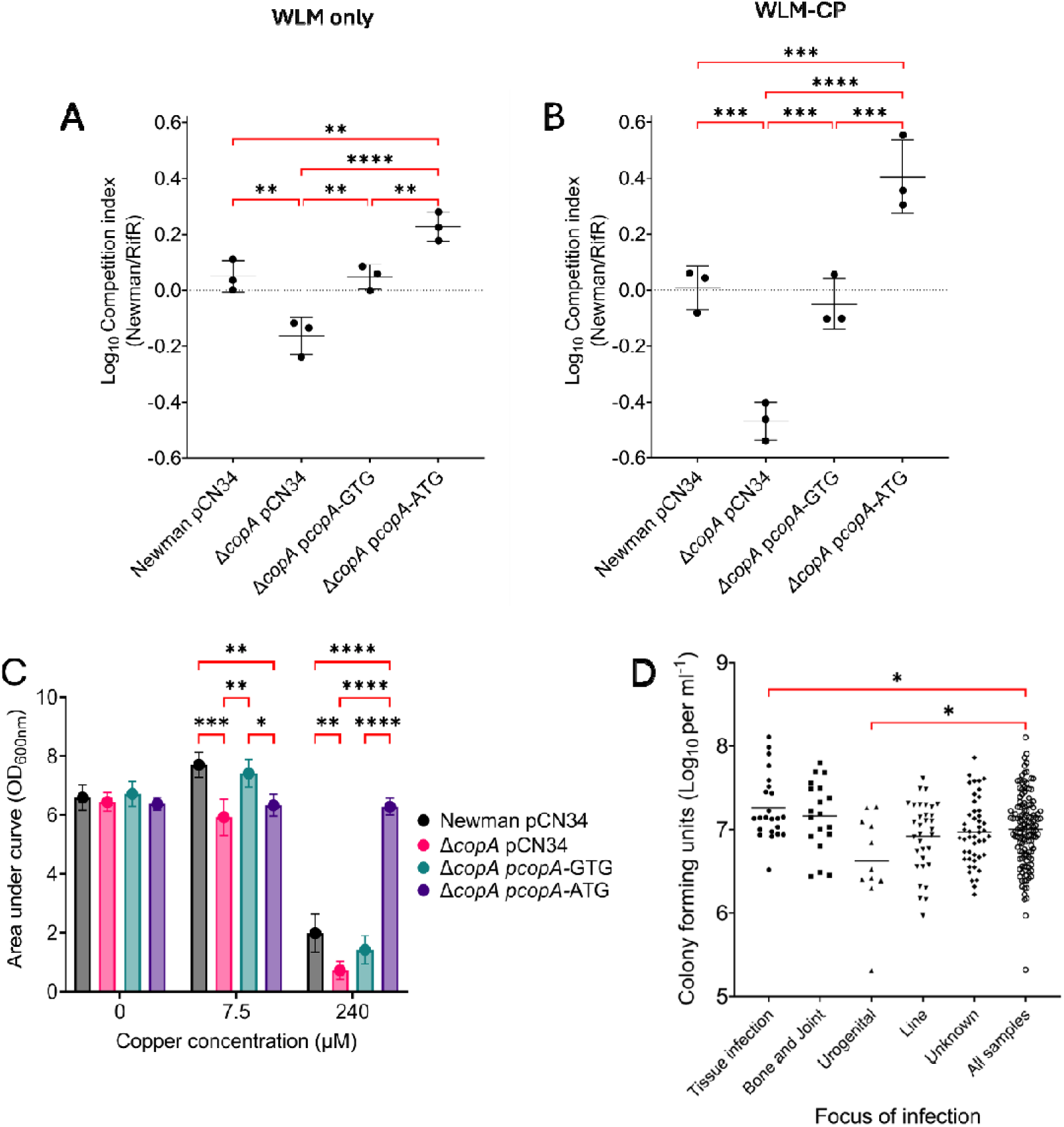
The *copA* SNP increases competitiveness of *S. aureus* in the presence of elevated copper, whilst reducing fitness in low copper environments. (A) Competition assay conducted in WLM. (B) Competition assay conducted in WLM-CP. (C) Growth assessment in CRPMI supplemented with low and high copper. (D) Comparison of CC22 clinical strains CFU outcomes in the polymicrobial WLM against the entry/focus of infection. Lines represent the mean of three independent experiments, error bars the standard deviation. Significance determined as *<0.05, **<0.01, ***<0.001, ****<0.0001 calculated using a one-way ANOVA with Sidak’s (A-B) or Dunnett’s (D) post-hoc test, and two-way ANOVA with Tukey’s multiple comparisons test (C)

As mentioned previously, the high number of clinical isolates expressing the GTG allele compared to ATG (124/134) suggests there must be environments where this confers a fitness benefit. Translation of genes with GTG open-reading frames provides organisms with greater translational control of metabolically costly proteins, with *S. aureus* using GTG for between 10-12% of its genes (45). As such we hypothesised that while the ATG *copA* allele conferred a fitness advantage in higher copper environments, such as WLM, the GTG codon likely confers an advantage in low copper environments, where the ATG driven production of CopA would export too much of this beneficial micronutrient. To test this we created a growth medium where we could more accurately control copper levels, by chelating the free metal ions from RPMI (CRPMI), and then add back in the other essential metal ions and a range of copper concentrations. In CRPMI devoid of copper, no difference in growth kinetics were observed across the strains (Fig 7c). However, upon the addition of a relatively low amount of copper (7.5μM; compared to normal serum copper levels, which are reported to range from 11.5 to 30 μM (46)) the wild type strain grew significantly better compared to the no-copper environment, whereas the growth of the *copA* mutant was impaired (Fig 7c). This demonstrates the benefit of this micronutrient for growth, but that intracellular control of its concentration by CopA is still required. Furthermore, the effect on growth of the mutant was complemented by p*copA*-GTG but not the p*copA*-ATG at 7.5μM copper (Fig 7c.), demonstrating that in this low copper environment, the GTG allele confers a greater fitness advantage than the ATG allele, presumably as the ATG allele removes too much copper. The converse was true when we increased the copper levels to those toxic for the bacteria (240μM), where the ATG allele was more beneficial for bacterial growth (Fig 7c), clearly highlighting how the relative benefit of each of these *copA* alleles crucially depends on the levels of copper in the environment.

Whilst the collection of strains used here were all isolated from bloodstream infections, the focus of infection before entry into the bloodstream varied, including skin and soft tissue infections, bone and joint infections, urogenital infections, and line infections. We sought to examine the clinical relevance of our findings by comparing the fitness of the strains in the WLM relative to their focus of infection and subsequent entry into the blood stream. We found that relative to the mean fitness of the whole collection, those from a skin and soft tissue infection origin were significantly more fit in the WLM, which most closely resembles this infection type. Moreover, eight of the 10 isolates with the ATG *copA* allele were from the skin and soft tissue entry point group, explaining the overall increase in fitness of this group of isolates (Table S3). Although the numbers are small, in the absence of any animal model in which to test this, this clinical data supports the concept that within-wound selection can occur in response to interkingdom interactions with copper a critical element to this.

## DISCUSSION

For *S. aureus* to colonise and persist at infection sites it must successfully compete with other microorganisms. These polymicrobial interactions can be synergistic or antagonistic in nature, with the outcome shaping the success of colonisation, establishment of infection and subsequent treatment and recovery (19,20,23,47). Colonisation of the host has long been associated with progression to infection, with asymptomatic *S. aureus* carriers being more prone to infections by this pathogen (48,49). However, how *S. aureus* competes with bacteria and fungi in these environments is poorly understood, with investigations being predominantly focussed on host-pathogen interactions with the bacteria in mono-culture (22). To address this, we applied a population-based approach to define the success of *S. aureus* establishment and persistence in a simple polymicrobial wound-like model with the opportunistic pathogens *C. albicans* and *P. aeruginosa*. In doing so we identified six proteins that contribute to the fitness of *S. aureus* in the presence of *P. aeruginosa* and *C. albicans* in the WLM, with the subsequent focus of this study being on the copper exporter CopA.

Our findings clearly demonstrate that the loss of CopA compromises the ability of *S. aureus* to persist in a wound like environment in the presence of *C. albicans* and *P. aeruginosa*. We found that *C. albicans* dependent release of copper from the host protein ceruloplasmin sensitised *S. aureus* to *P. aeruginosa* produced redox active antimicrobials pyocyanin and KCN (Fig 8A). In particular, we identified a single point mutation (1G>A) that enhanced translation of the CopA exporter, enabling robust control of copper accumulation in *S. aureus*. The reduction in cytoplasmic copper accumulation with *copA* 1G>A resulted in improved fitness in our polymicrobial biofilm model in both clinical and laboratory strains and was associated with reduced susceptibility to pyocyanin and KCN, with the enhanced control of copper mitigating the toxicity of these *P. aeruginosa* metabolites (Figure 8B). The presence of the ATG *copA* allele and its association with clinical skin and soft tissue infections may represent infection site specific evolution of these *S. aureus* isolates, given the fitness benefit it confers in this high copper environment.

**Figure 8.**
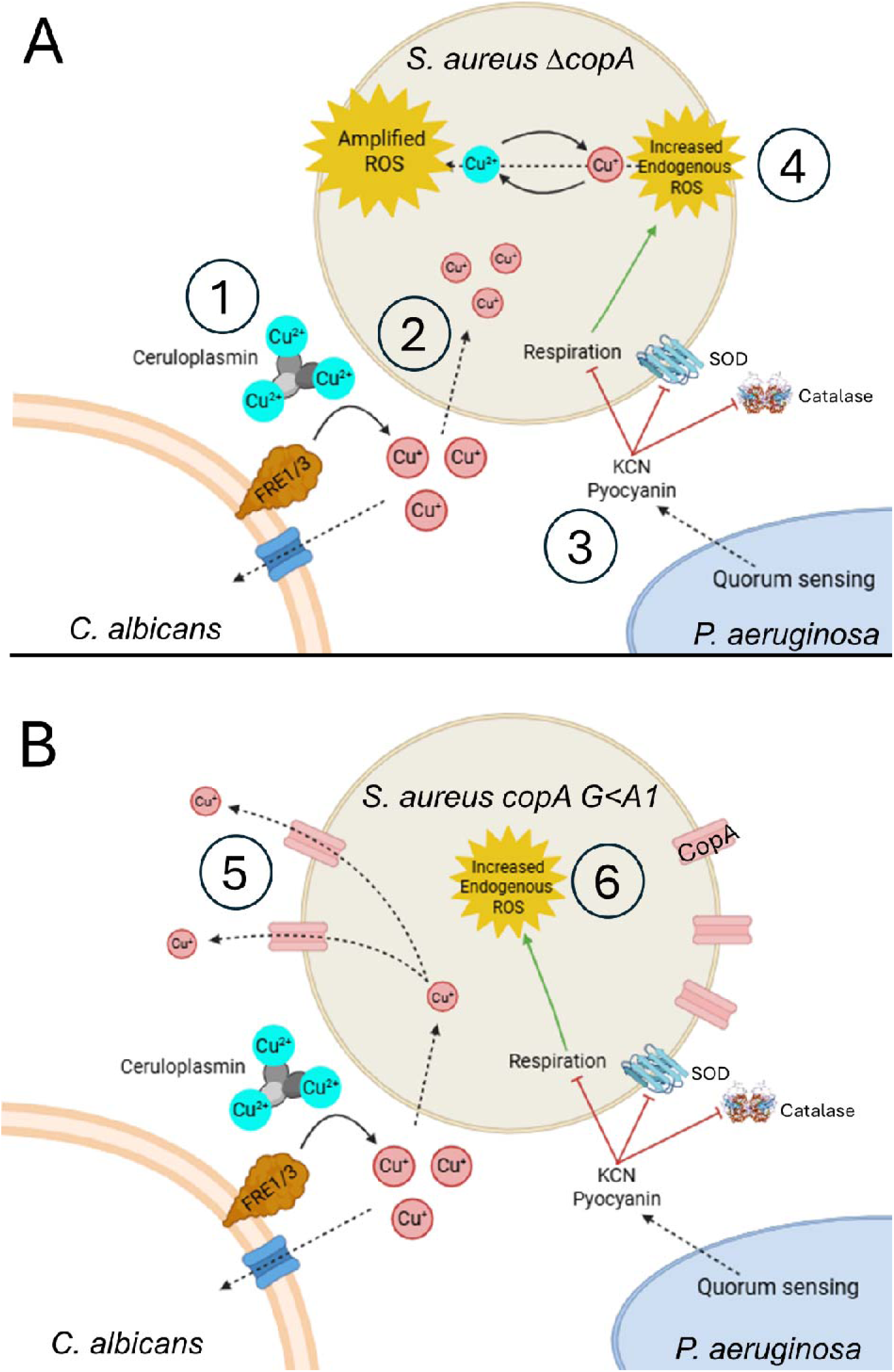
Graphical summary of our proposed mechanism surrounding copper accumulation and *S. aureus* fitness in polymicrobial wound environments. Mechanism in the absence of a functional copper exporter (A), and the proposed role of the *copA* 1G>A allele in the polymicrobial WLM (B). 1) Copper bound to host produced ceruloplasmin is reduced, liberating Cu^+^. 2) Free copper is imported by *C. albicans* and *S. aureus*, in the absence of a functional copper exporter Cu^+^ accumulates in the *S. aureus* cytoplasm. 3) *P. aeruginosa* quorum-sensing circuit induces production of redox active antimicrobials pyocyanin and cyanide. Both inhibit respiration leading to increased production of ROS, and inactivate superoxide dismutase/catalase, compromising the ability of *S. aureus* to neutralise accumulating ROS. 4) The presence of copper amplifies the increased endogenous ROS though Fenton-like chemistry, catalysing production superoxide and hydroxyl radicals from oxygen and H_2_O_2_ respectively. In combination *C. albicans* derived copper combined with the action of *P. aeruginosa* antimicrobials leads to amplification of ROS in *S. aureus* unable to export copper. As a result, these strains are unable to maintain viability and exhibit reduced persistence in polymicrobial communities. 5) The *copA* G>A1 identified as demonstrating enhanced competition in this community demonstrates decreased cytoplasmic copper accumulation due to an increase in the abundance of CopA. 6) Removal of copper reduces the incidence of copper mediated ROS generation enabling strains encoding *copA 1*G<A to successfully maintain itself in the presence of *C. albicans* and *P. aeruginosa*.

That bacteria such as *S. aureus* have evolved complex copper uptake and efflux systems demonstrates the importance to them of balancing the intracellular homeostasis of this element. However, the role of copper during infection is complex, with only 5% of copper in plasma being in a freely available form and the remainder bound by ceruloplasmin (50). Ceruloplasmin concentrations vary depending on the body site, with a normal serum concentration of 20-50 mg/dL (51). Infection is commonly believed to lead to an increase in ceruloplasmin, although the exact level of increase is highly variable (32–34). Neutrophils and macrophages release copper from ceruloplasmin and pump copper ions into phagosomes to promote bacterial killing (32,33). While the effect we report here on *S. aureus* fitness is due to polymicrobial interactions, given that the innate immune system serves as a source of both copper and redox antimicrobials, the likely scenario is that both the host innate immune system and microbial population at infection sites will work cooperatively to protect against *S. aureus* infection.

Our findings are in some ways at odds with what has been suggested previously, where beneficial interactions between *C. albicans* and *S. aureus* have been reported under biofilm conditions. For example, *C. albicans* polysaccharide production results in formation of a polymicrobial biofilms, which protects both *S. aureus* and *C. albicans* from the host innate immune system and antimicrobials (20,21). The detection of hydrolysed *S. aureus* peptidoglycan by *C. albicans* triggers hyphae formation that contributes to polymicrobial biofilm production (52), whereby *S. aureus* attaches to these hyphae via the Als3p protein (53,54). The formation of hypoxic microenvironments within these extensive biofilm structures has been shown to enhance *S. aureus* toxin production (47), which will likely cause further host tissue damage and can promote further *C. albicans* hyphae development (55). Instead, here we report an antagonist interaction where copper is used by *C. albicans* as a toxic element to enhance *S. aureus* killing. It is however worth considering that this only occurred in the presence of a third microorganism with potent redox reactive antimicrobial activity: *P. aeruginosa.* In an otherwise low copper environment, the provision of free copper by *C. albicans* may represent a benefit to *S. aureus* given its ability to enhance growth. As such, the likely cost-benefit of this copper-based interaction is likely to depend on the presence of such antimicrobial agents, be they of microbial or innate immune system origin.

In addition to *copA*, five other genes were identified to play a role in the fitness of *S. aureus* in a polymicrobial wound model: *ftsK*, *rluD*, *ribBA*, *cntB* and a gene with the locus tag SAUSA300_2579. The DNA translocase FtsK and the pseudouridine synthase RluD are involved with chromosomal segregation and ribosome assembly (56,57). As such, the reduced fitness phenotype may be due to reduced growth rates when compared to the isogenic wildtype strain; however, why such mutations might emerge in natural populations of *S. aureus* remains to be determined. RibBA is required for *de novo* synthesis of riboflavin and flavin nucleotides, where riboflavin has antioxidant properties, so perhaps its role here is protective against the redox active antimicrobial compounds produced by *P. aeruginos*a (58). CntB is an ABC transporter, which forms part of the staphylopine transport complex. Staphylopine is used to scavenge and transport multiple metals including nickel, cobalt, zinc, copper and manganese (59). In contrast to CopA, which exports copper, loss of CntB impairs the ability of *S. aureus* to acquire zinc and other essential metal ions (60), likely represents nutritional competition with both *C. albicans* and *P. aeruginosa.* However, the identification of another metal ion transport system highlights the importance of metal ion homeostasis in facilitating *S. aureus* competition in polymicrobial communities. The potential contribution of the protein encoded by SAUSA300_2579, which is an amidase-domain containing protein, is currently unclear.

The importance of transition metals as cofactors for biological processes is such that the vertebrate immune system has evolved to control access to them during infections. This process of nutritional immunity can either starve pathogens of essential metals or accumulate them in excess during inflammation to intoxicate the pathogen. Copper appears to fall into the latter category; and while the role of copper in host protection during infection is incompletely understood, this study brings an entirely new dimension to our understanding of this. In addition to the host using copper to control infections through innate immune pathways, here we show that competing pathogens from different kingdoms of life can work together to take this copper and use it to specifically target a fellow invading pathogen. While this new understanding opens up the tantalising prospect of the use of single chemical element as a means of therapeutic intervention, *S. aureus* has already found ways around this through its mutation of *copA*. This adaptation is however likely shortsighted in outcome, given the potential fitness costs that increasing copper efflux would have once the infection has cleared and the bacteria has transitioned back to commensalism.

In summary, this study demonstrates how a single chemical element can be central to interkingdom interactions during infection processes, which in turn emphasises how the contribution of the infection ecosystem in its entirety should be considered to gain a more complete understanding of *S. aureus* colonisation and persistence at infection sites.

## MATERIALS AND METHODS

### Bacterial strains, plasmids and oligonucleotides

Bacterial strains and plasmids used in this study are listed in table 1. Clinical CC22 isolates used for GWAS study are reported in (13). *S. aureus* strains were routinely maintained on TSA, *P. aeruginosa* and *E. coli* on LBA, and *C. albicans* on SDA. When required, antibiotics were added at concentrations of 10 µg mL^-1^ for erythromycin and chloramphenicol, 90 µg mL^-1^ kanamycin, 100 µg mL^-1^ carbenicillin, 125 µg mL^-1^ tetracycline and 10 µg mL^-1^ nystatin.

**Table 1.**
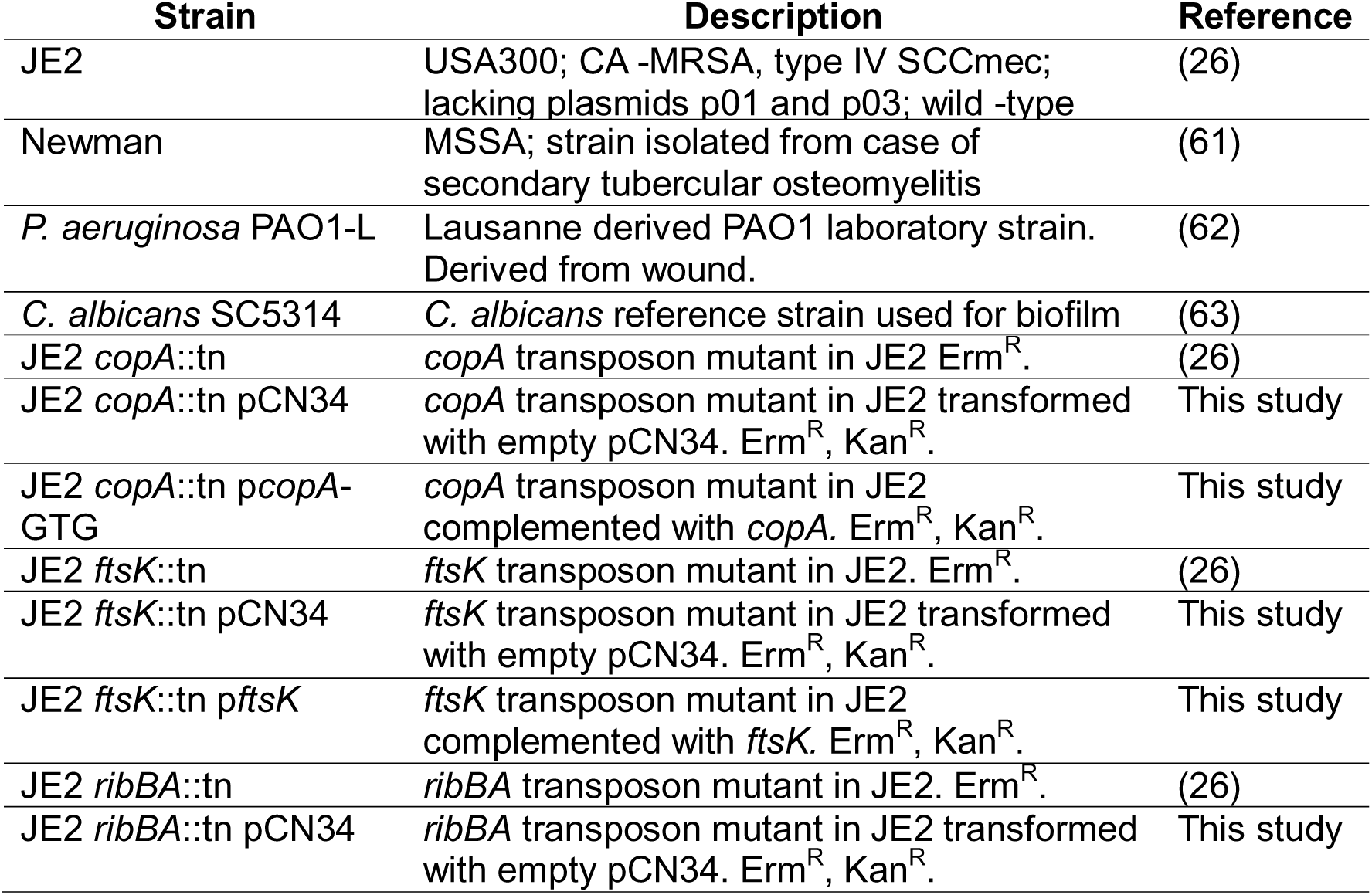
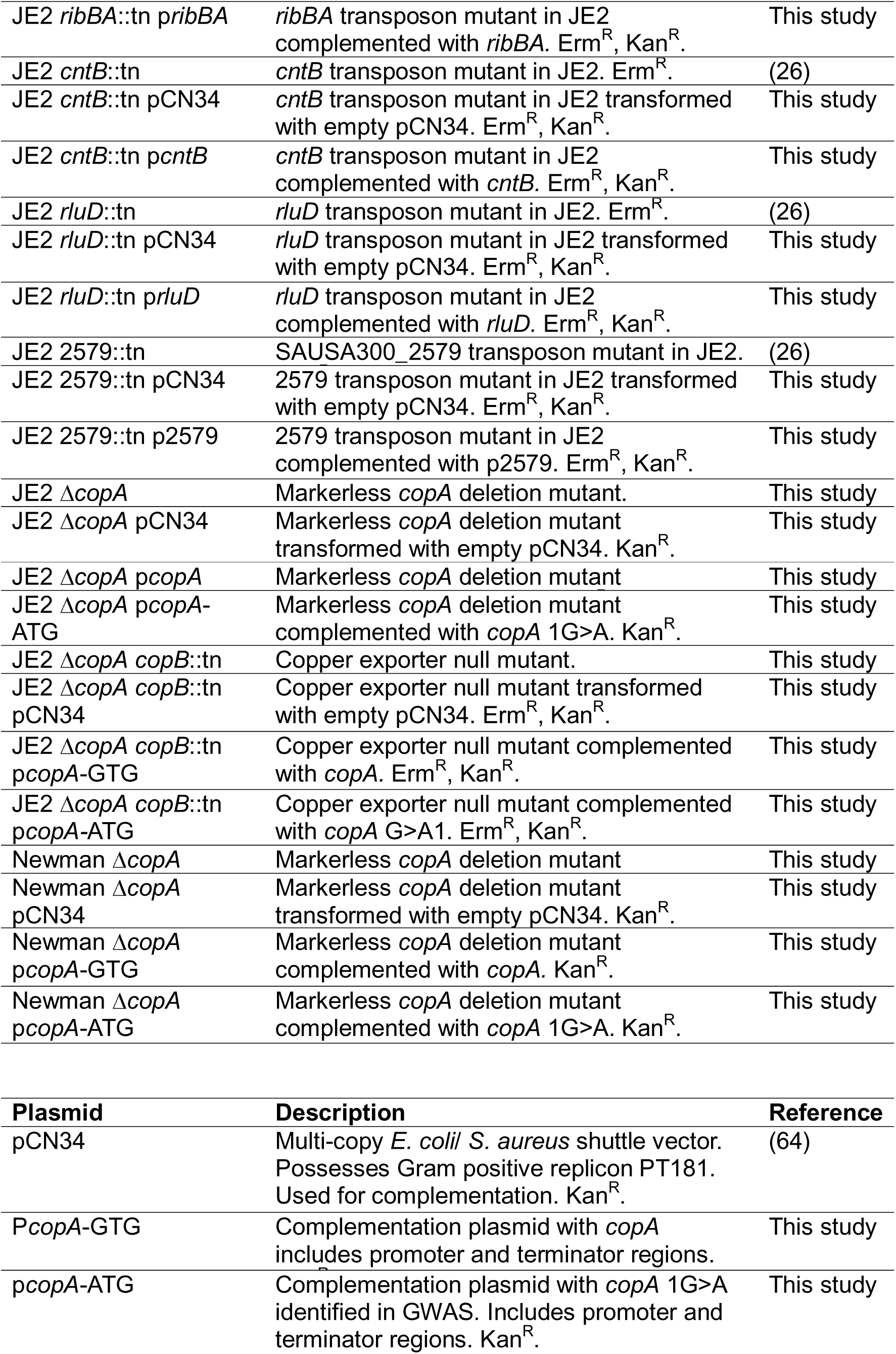
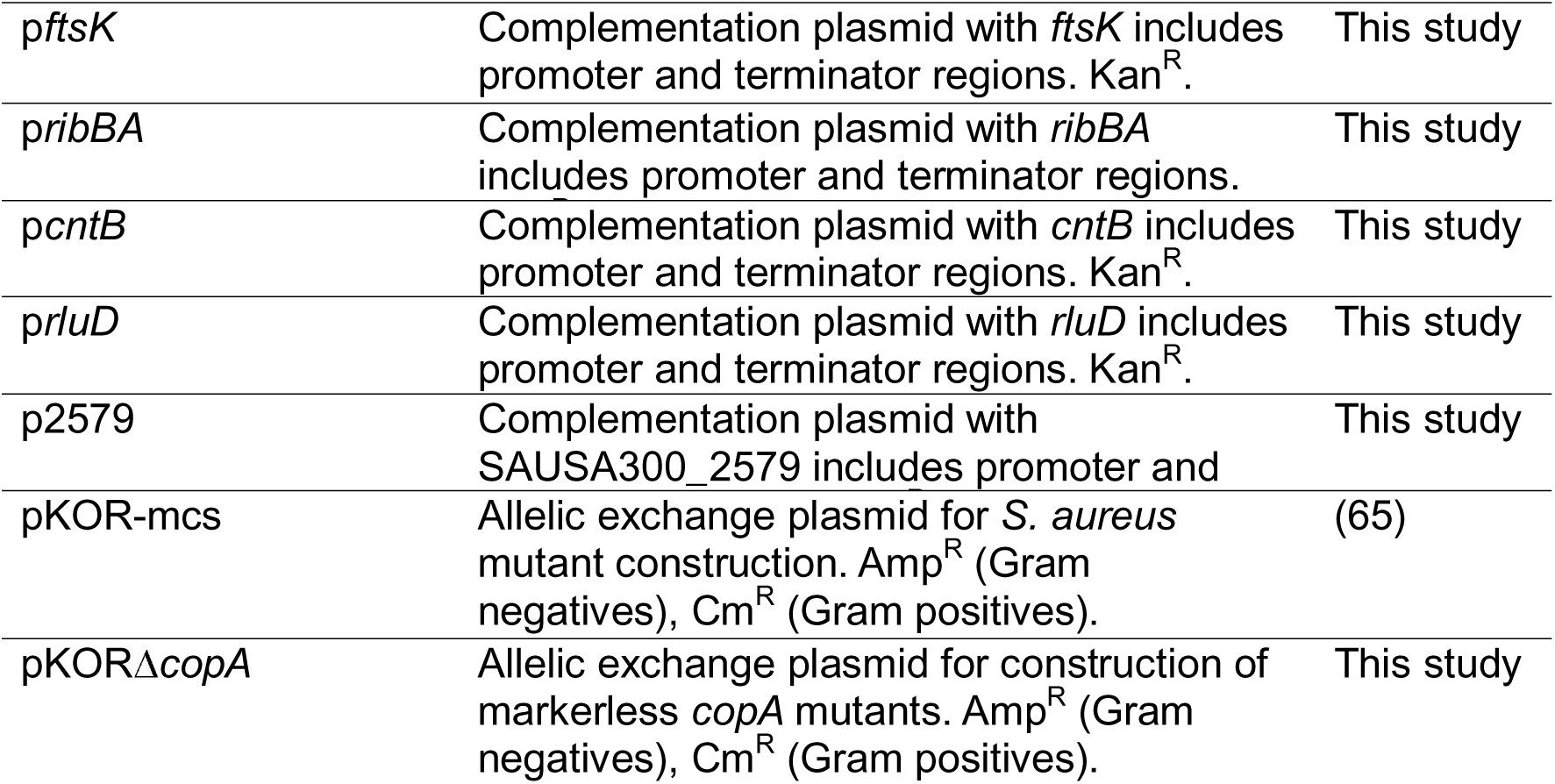
Strains and plasmids used in this study.

### Wound-like media polymicrobial model

For the initial GWAS screen WLM consisted of 50% bovine plasma (VWR), 5% horse blood (Oxoid) and 45% bolton broth (Oxoid) as previously reported by Dalton and colleagues (66). For validation of transposon mutants and thereafter, media was altered to 50% human plasma (Clinisciences), 5% human blood (Clinisciences) and 45% Bolton broth to make media specific to human wound infections. *S. aureus*, *C. albicans* and *P. aeruginosa* strains were inoculated into 200 µL of WLM at 1×10^3^ CFU mL^-1^ in 2 mL screwcap tubes and incubated statically at 37^°^C for 1-7 days. At endpoint two ceramic zirconium beads (2.6mm) were added to each tube alongside 800 µL of PBS. Biofilms were then disaggregated using tissue homogenization (MP Biomedicals), 10-fold serially diluted in PBS, and plated on selective media for enumeration. *S. aureus* and *P. aeruginosa* were selectively plated on mannitol salt and *Pseudomonas* isolation agar respectively, with nystatin added to prevent *C. albicans* recovery. For recovery of *C. albicans*, biofilms were plated on Sabouraed dextrose agar with tetracycline to inhibit *S. aureus* and *P. aeruginosa* recovery. For the GWAS, biofilms were quantified at the 3-day timepoint only. For continuous experiments at least three biofilms were sacrificed per timepoint.

### Genome Wide Association Study

Genome-wide association mapping was conducted using a generalised linear model as previously described (67). *S. aureus* CFU recovery in a polymicrobial community was assigned as the quantitative response variable. We accounted for bacterial population substructure by adding to the regression model the first two component from a principal component decomposition of SNP data for the CC22 clinical samples, which accounted for 32% of the total variance. In total, 2066 unique SNPs were analysed. The p-values reported in table S1 are not corrected for multiple comparisons.

### DNA manipulations, mutant construction and complementation

Genomic DNA isolation was performed using the GenElute gDNA purification kit (Sigma). Plasmid minipreps and PCR purifications performed with Qiaprep spin mini kit (Qiagen). PCR products for cloning applications amplified with Phusion polymerase (Thermo). Restriction digest, ligations and transformation performed according to standard protocols. Oligonucleotides (Sigma) used in this study can be found in table 2.

**Table 2.**
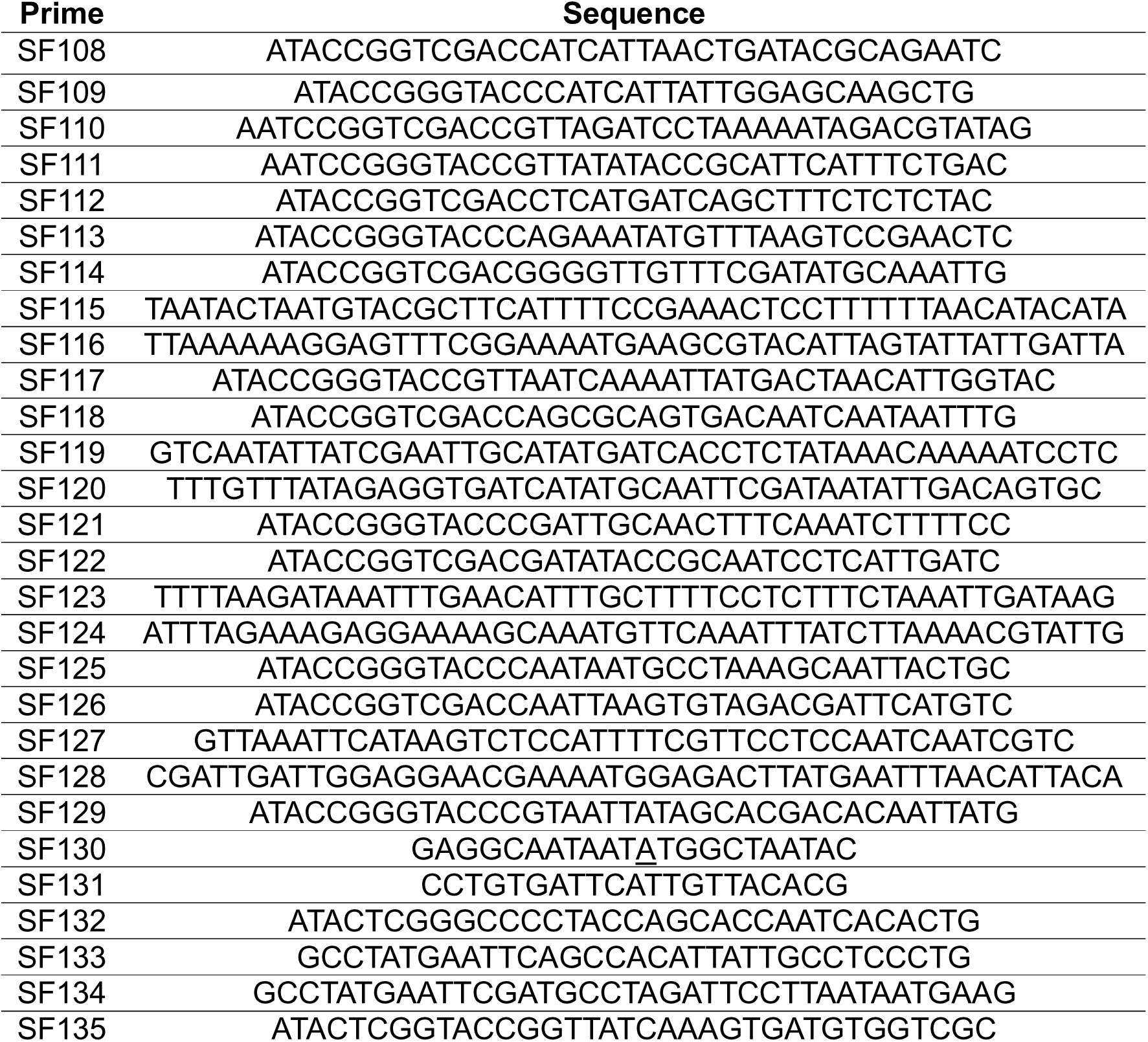
Oligonucleotides used in this study. Prime Sequence.

Markerless *copA* mutants were constructed in *S. aureus* using the allelic exchange vector pKOR-mcs (65). Two DNA fragments consisting of the upstream 1202bp and downstream 1092bp were PCR amplified using primer pairs SF132/SF133 and SF134/SF135 respectively. These products were fused by overlap extension PCR and ligated into pKOR-mcs using the ApaI/KpnI restriction sites introduced on primers SF132 and SF135, forming suicide plasmid pKORΔ*copA*. Ligation mixtures were transformed into chemically competent *E. coli* IM08B, miniprepped, and insertion confirmed by PCR and Sanger sequencing. After electroporation into *S. aureus* JE2, JE2 *copB*::tn and Newman, allelic exchange was induced by temperature shift and plating on anhydrotetracycline according to procedures outlined in (65,68).

Complementation of validated GWAS targets performed using the pCN34 plasmid (64). The *copA* gene including the native promoter was amplified from *S. aureus* JE2 using primer pair SF108/109; *ftsK* with SF110/111; and NE1353 with SF112/113. For genes in an operon, the open reading frame corresponding to the target gene and the promoter sequence upstream of the first gene in the operon were fused together by overlap extension PCR using primers SF114-117 for *camS*; SF118-121 for *ribBA*; SF122-125 for *cntB*; and SF126-129 for *rluD*. These products were then ligated into pCN34 using SalI/KpnI restriction sites forming complementation plasmids p*copA*-GTG, p*ftsK,* p*camS,* p*ribBA,* p*rluD and* p*NE1353.* Ligations were directly transformed into *E. coli* IM08B (69), purified and then electroporated into the required *S. aureus* strains. For generation of p*copA*-ATG with the 1G>A mutation identified in the GWAS site-directed mutagenesis was performed. The p*copA*-GTG plasmid was used as a template for primer pair SF130/SF131, with the 1G>A mutation introduced in the middle of primer SF130. This PCR product was PCR purified and simultaneously treated with DpnI and T4 polynucleotide kinase to remove plasmid template and phosphorylate the newly generated p*copA* G>A1 amplicon. The fragment was then ligated, transformed into *E. coli* IM08B, and subsequently electroporated into the required *S. aureus* strains.

### Intracellular Copper quantification

WLM biofilms were established in the presence and absence of 20 µM human ceruloplasmin and plated for CFUs. Cells from six biofilms were pooled and concentrated into 200 µL of HPLC-grade water. Intracellular contents of *S. aureus* cells were released through treatment with 50 µg mL^-1^ of lysostaphin for 2 H followed by measurement of copper using a Copper assay kit (Sigma) according to manufacturer’s instructions. Values obtained were normalised to total CFU count and normalised to copper contained in 1×10^7^ *S. aureus*.

### Relative Fitness Assay

Relative fitness assays were conducted as reported in (70)*.S. aureus* isolates were cultured overnight in TSB. Competitions were established in WLM with and without exogenous ceruloplasmin at a concentration of 20 µg mL^-1^. The WLM and WLM-CP were inoculated with 5×10^2^ of the rifampicin resistant marker strain (RIF1) and the test strain with initial numbers confirmed by plating. *P. aeruginosa* and *C. albicans* were inoculated when required as detail in the WLM model. Strains were competed as single-species or polymicrobial biofilms at 37°C for 3 days. Final cell numbers were enumerated by plating serial dilutions on TSA (total cell count) and TSA with 1 µg mL^-1^ rifampicin (marker strain count). The fitness of the strain is defined as a measure of reproduction success of the population using the below equation. Each competition assay was performed in triplicate with three independent experiments.

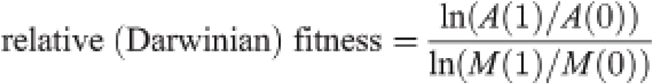

Where: *A*(0), estimated density of test strain at time 0; *M*(0), estimated density of marker strain at time 0; *A*(1), estimated density of test strain at time 1 d; *M*(1), estimated density of marker strain at time 1 d; ln, natural logarithm (logarithm to the base *e*).

### Redox-active antimicrobial killing/sensitivity assays

Prior to killing assays strains were cultured for 3 days as polymicrobial biofilms in the WLM with or without 20 µM ceruloplasmin. For H_2_O_2_ killing assays the number of *S. aureus* contained within the community was normalised to 1×10^6^ in PBS. *S. aureus* was then exposed to 20 mM H_2_O_2_ for 3 H at 37^°^C. For pyocyanin (Sigma), KCN (Sigma) and HQNO assays, *S. aureus* was normalised to 1×10^6^ in TSB. Strains were exposed to 0.2 mM pyocyanin, 1 mM KCN and 0.2 mM HQNO for 6 H at 37°C. After treatment strains were serially diluted in Dey-Engley neutralisation broth and plated for CFU recovery on mannitol salt agar with nystatin (10 µg mL^-1^). Thiourea protection assays were performed similarly, however prior to the final dilution *S. aureus* was incubated with 150 mM thiourea in PBS for 15 minutes before washing and treatment with H_2_O_2_, pyocyanin, KCN or HQNO.

### Metal ion removal from RPMI

Metal ions from RPMI 1640 medium (thermofisher: 11875093) were removed with Chelex-100 (Sigma) according to (71) with modifications. RPMI was amended with 1% casamino acids. Chelex-100 was mixed with RPMI (5% w/v) for 1 H with shaking and filter sterilised. The following essential ions were re-introduced: ZnCl_2_ (25 μM), CaCl_2_ (100 μM), MgCl_2_ (1 mM), MnCl_2_ (25 μM) and FeCl_3_ (25 μM). CuCl_2_ was added at the reported concentrations when required. Strains were then cultured in 96-well plate format using this media at 37°C with shaking. Readings (OD_600nm_) were kinetically tracked for 16 H for each strain and copper concentration using a Tecan infinite pro 200 plate reader. Resulting growth kinetics were analysed on GraphPad Prism (V10.4.1), using the area under the curve function.

## ACKNOWLEDGEMENTS

All authors acknowledge the provision of strain by the Network on Antimicrobial Resistance in *Staphylococcus aureus* (NARSA) Program: under NIAID/NIH Contract number HHSN272200700055C. We acknowledge financial support from Research Ireland Frontiers for Future programme grant (reference: 21/FFP-A/9704) and a Wellcome Trust Discovery Award (ref: 227883/23/Z). We thank Professor Miguel Camara (University of Nottingham) and Dr Jerry Reen for providing *P. aeruginosa* PAO1-L and *C. albicans* SC5314 respectively. Allelic exchange plasmid pKOR-mcs was a gift from Suzan Rooijakkers (Addgene plasmid #181755; http://n2t.net/addgene:181755; RRID: Addgene_181755). We also thank members of both the Cork and Bristol team for helpful discussion of this work.

## DATA AVAILIBILITY STATEMENT

The authors confirm that the data supporting the findings of this study are available with the article and supplementary material. Bacterial strains developed as part of this study are available upon request.

## REFERENCES

1. Antimicrobial Resistance Collaborators. Global burden of bacterial antimicrobial resistance in 2019: a systematic analysis. Lancet Lond Engl. 2022 Feb 12;399(10325):629–55.

2. Gordon RJ, Lowy FD. Pathogenesis of methicillin-resistant Staphylococcus aureus infection. Clin Infect Dis Off Publ Infect Dis Soc Am. 2008 June 1;46 Suppl 5(Suppl 5):S350–359.

3. Krismer B, Weidenmaier C, Zipperer A, Peschel A. The commensal lifestyle of Staphylococcus aureus and its interactions with the nasal microbiota. Nat Rev Microbiol. 2017 Oct 12;15(11):675–87.

4. Wertheim HFL, Melles DC, Vos MC, van Leeuwen W, van Belkum A, Verbrugh HA, et al. The role of nasal carriage in Staphylococcus aureus infections. Lancet Infect Dis. 2005 Dec;5(12):751–62.

5. Queen D, Harding K. Estimating the cost of wounds both nationally and regionally within the top 10 highest spenders. Int Wound J. 2024 Feb;21(2):e14709.

6. Dowd SE, Sun Y, Secor PR, Rhoads DD, Wolcott BM, James GA, et al. Survey of bacterial diversity in chronic wounds using pyrosequencing, DGGE, and full ribosome shotgun sequencing. BMC Microbiol. 2008 Mar 6;8:43.

7. Bessa LJ, Fazii P, Di Giulio M, Cellini L. Bacterial isolates from infected wounds and their antibiotic susceptibility pattern: some remarks about wound infection. Int Wound J. 2015 Feb;12(1):47–52.

8. Puca V, Marulli RZ, Grande R, Vitale I, Niro A, Molinaro G, et al. Microbial Species Isolated from Infected Wounds and Antimicrobial Resistance Analysis: Data Emerging from a Three-Years Retrospective Study. Antibiot Basel Switz. 2021 Sept 24;10(10):1162.

9. Ge Y, Wang Q. Current research on fungi in chronic wounds. Front Mol Biosci. 2022;9:1057766.

10. Douglas EJA, Palk N, Brignoli T, Altwiley D, Boura M, Laabei M, et al. Extensive remodelling of the cell wall during the development of Staphylococcus aureus bacteraemia. eLife. 2023 July 4;12:RP87026.

11. Laabei M, Uhlemann AC, Lowy FD, Austin ED, Yokoyama M, Ouadi K, et al. Evolutionary Trade-Offs Underlie the Multi-faceted Virulence of Staphylococcus aureus. PLoS Biol. 2015;13(9):e1002229.

12. Laabei M, Recker M, Rudkin JK, Aldeljawi M, Gulay Z, Sloan TJ, et al. Predicting the virulence of MRSA from its genome sequence. Genome Res. 2014 May;24(5):839–49.

13. Recker M, Laabei M, Toleman MS, Reuter S, Saunderson RB, Blane B, et al. Clonal differences in Staphylococcus aureus bacteraemia-associated mortality. Nat Microbiol. 2017 Oct;2(10):1381–8.

14. Altwiley D, Brignoli T, Edwards A, Recker M, Lee JC, Massey RC. A functional menadione biosynthesis pathway is required for capsule production by Staphylococcus aureus. Microbiol Read Engl. 2021 Nov;167(11):001108.

15. Stevens EJ, Morse DJ, Bonini D, Duggan S, Brignoli T, Recker M, et al. Targeted control of pneumolysin production by a mobile genetic element in Streptococcus pneumoniae. Microb Genomics. 2022 Apr;8(4):000784.

16. Douglas EJA, Duggan S, Brignoli T, Massey RC. The MpsB protein contributes to both the toxicity and immune evasion capacity of Staphylococcus aureus. Microbiol Read Engl. 2021 Oct;167(10):001096.

17. Murray JL, Connell JL, Stacy A, Turner KH, Whiteley M. Mechanisms of synergy in polymicrobial infections. J Microbiol Seoul Korea. 2014 Mar;52(3):188–99.

18. Hotterbeekx A, Kumar-Singh S, Goossens H, Malhotra-Kumar S. In vivo and In vitro Interactions between Pseudomonas aeruginosa and Staphylococcus spp. Front Cell Infect Microbiol. 2017;7:106.

19. Nair N, Biswas R, Götz F, Biswas L. Impact of Staphylococcus aureus on pathogenesis in polymicrobial infections. Infect Immun. 2014 June;82(6):2162–9.

20. Kong EF, Tsui C, Kucharíková S, Andes D, Van Dijck P, Jabra-Rizk MA. Commensal Protection of Staphylococcus aureus against Antimicrobials by Candida albicans Biofilm Matrix. mBio. 2016 Oct 11;7(5):e01365–16.

21. Harriott MM, Noverr MC. Candida albicans and Staphylococcus aureus form polymicrobial biofilms: effects on antimicrobial resistance. Antimicrob Agents Chemother. 2009 Sept;53(9):3914–22.

22. Peters BM, Jabra-Rizk MA, O’May GA, Costerton JW, Shirtliff ME. Polymicrobial interactions: impact on pathogenesis and human disease. Clin Microbiol Rev. 2012 Jan;25(1):193–213.

23. Keim K, Bhattacharya M, Crosby HA, Jenul C, Mills K, Schurr M, et al. Polymicrobial interactions between Staphylococcus aureus and Pseudomonas aeruginosa promote biofilm formation and persistence in chronic wound infections. BioRxiv Prepr Serv Biol. 2024 Nov 5;2024.11.04.621402.

24. Vestweber PK, Wächter J, Planz V, Jung N, Windbergs M. The interplay of Pseudomonas aeruginosa and Staphylococcus aureus in dual-species biofilms impacts development, antibiotic resistance and virulence of biofilms in in vitro wound infection models. PloS One. 2024;19(5):e0304491.

25. DeLeon S, Clinton A, Fowler H, Everett J, Horswill AR, Rumbaugh KP. Synergistic interactions of Pseudomonas aeruginosa and Staphylococcus aureus in an in vitro wound model. Infect Immun. 2014 Nov;82(11):4718–28.

26. Fey PD, Endres JL, Yajjala VK, Widhelm TJ, Boissy RJ, Bose JL, et al. A genetic resource for rapid and comprehensive phenotype screening of nonessential Staphylococcus aureus genes. mBio. 2013 Feb 12;4(1):e00537–00512.

27. Zhao L, Li H, Liu Z, Wang Z, Xu D, Zhang J, et al. Copper ions induces ferroptosis in Staphylococcus aureus and promotes healing of MRSA-induced wound infections. Microbiol Res. 2025 July;296:128122.

28. Baker J, Sitthisak S, Sengupta M, Johnson M, Jayaswal RK, Morrissey JA. Copper stress induces a global stress response in Staphylococcus aureus and represses sae and agr expression and biofilm formation. Appl Environ Microbiol. 2010 Jan;76(1):150–60.

29. Purves J, Thomas J, Riboldi GP, Zapotoczna M, Tarrant E, Andrew PW, et al. A horizontally gene transferred copper resistance locus confers hyper-resistance to antibacterial copper toxicity and enables survival of community acquired methicillin resistant Staphylococcus aureus USA300 in macrophages. Environ Microbiol. 2018 Apr;20(4):1576–89.

30. Sitthisak S, Knutsson L, Webb JW, Jayaswal RK. Molecular characterization of the copper transport system in Staphylococcus aureus. Microbiol Read Engl. 2007 Dec;153(Pt 12):4274–83.

31. Natesha RK, Natesha R, Victory D, Barnwell SP, Hoover EL. A prognostic role for ceruloplasmin in the diagnosis of indolent and recurrent inflammation. J Natl Med Assoc. 1992 Sept;84(9):781–4.

32. White C, Lee J, Kambe T, Fritsche K, Petris MJ. A role for the ATP7A copper-transporting ATPase in macrophage bactericidal activity. J Biol Chem. 2009 Dec 4;284(49):33949–56.

33. Achard MES, Stafford SL, Bokil NJ, Chartres J, Bernhardt PV, Schembri MA, et al. Copper redistribution in murine macrophages in response to Salmonella infection. Biochem J. 2012 May 15;444(1):51–7.

34. Besold AN, Shanbhag V, Petris MJ, Culotta VC. Ceruloplasmin as a source of Cu for a fungal pathogen. J Inorg Biochem. 2021 June;219:111424.

35. Pham AN, Xing G, Miller CJ, Waite TD. Fenton-like copper redox chemistry revisited: Hydrogen peroxide and superoxide mediation of copper-catalyzed oxidant production. J Catal. 2013 May;301:54–64.

36. Li H, Zhou X, Huang Y, Liao B, Cheng L, Ren B. Reactive Oxygen Species in Pathogen Clearance: The Killing Mechanisms, the Adaption Response, and the Side Effects. Front Microbiol. 2020;11:622534.

37. Beavers WN, Skaar EP. Neutrophil-generated oxidative stress and protein damage in*Staphylococcus aureus*. Napier B, editor. Pathog Dis. 2016 Aug;74(6):ftw060.

38. Noto MJ, Burns WJ, Beavers WN, Skaar EP. Mechanisms of Pyocyanin Toxicity and Genetic Determinants of Resistance in Staphylococcus aureus. J Bacteriol. 2017 Sept 1;199(17):e00221–17.

39. Létoffé S, Wu Y, Darch SE, Beloin C, Whiteley M, Touqui L, et al. Pseudomonas aeruginosa Production of Hydrogen Cyanide Leads to Airborne Control of Staphylococcus aureus Growth in Biofilm and In Vivo Lung Environments. mBio. 2022 Oct 26;13(5):e0215422.

40. Hoffman LR, Déziel E, D’Argenio DA, Lépine F, Emerson J, McNamara S, et al. Selection for Staphylococcus aureus small-colony variants due to growth in the presence of Pseudomonas aeruginosa. Proc Natl Acad Sci U S A. 2006 Dec 26;103(52):19890–5.

41. O’Malley YQ, Reszka KJ, Rasmussen GT, Abdalla MY, Denning GM, Britigan BE. The Pseudomonas secretory product pyocyanin inhibits catalase activity in human lung epithelial cells. Am J Physiol Lung Cell Mol Physiol. 2003 Nov;285(5):L1077–1086.

42. Hassett DJ, Schweizer HP, Ohman DE. Pseudomonas aeruginosa sodA and sodB mutants defective in manganese- and iron-cofactored superoxide dismutase activity demonstrate the importance of the iron-cofactored form in aerobic metabolism. J Bacteriol. 1995 Nov;177(22):6330–7.

43. Ogura Y, Yamazaki I. Steady-state kinetics of the catalase reaction in the presence of cyanide. J Biochem (Tokyo). 1983 Aug;94(2):403–8.

44. Shearer J, Fitch SB, Kaminsky W, Benedict J, Scarrow RC, Kovacs JA. How does cyanide inhibit superoxide reductase? Insight from synthetic FeIIIN4S model complexes. Proc Natl Acad Sci U S A. 2003 Apr 1;100(7):3671–6.

45. Cherrak Y, Salazar MA, Näpflin N, Malfertheiner L, Herzog MKM, Schubert C, et al. Non-canonical start codons confer context-dependent advantages in carbohydrate utilization for commensal E. coli in the murine gut. Nat Microbiol. 2024 Oct;9(10):2696–709.

46. Andrew D, Gail R, Morag B, Kishor R. Recommended reference intervals for copper and zinc in serum using the US National Health and Nutrition Examination surveys (NHANES) data. Clin Chim Acta Int J Clin Chem. 2023 June 1;546:117397.

47. Todd OA, Fidel PL, Harro JM, Hilliard JJ, Tkaczyk C, Sellman BR, et al. Candida albicans Augments Staphylococcus aureus Virulence by Engaging the Staphylococcal agr Quorum Sensing System. mBio. 2019 June 4;10(3):e00910–19.

48. Wertheim HFL, Vos MC, Ott A, van Belkum A, Voss A, Kluytmans JAJW, et al. Risk and outcome of nosocomial Staphylococcus aureus bacteraemia in nasal carriers versus non-carriers. Lancet Lond Engl. 2004 Aug 21;364(9435):703–5.

49. Dinh KM, Kaspersen KA, Boldsen JK, Ellermann-Eriksen S, Ostrowski SR, Aagaard B, et al. Evaluating infection risk associated with Staphylococcus aureus nasal carriage in blood donors: a prospective multicentre study in Denmark. Clin Microbiol Infect Off Publ Eur Soc Clin Microbiol Infect Dis. 2025 July;31(7):1180–6.

50. Catalani S, Paganelli M, Gilberti ME, Rozzini L, Lanfranchi F, Padovani A, et al. Free copper in serum: An analytical challenge and its possible applications. J Trace Elem Med Biol Organ Soc Miner Trace Elem GMS. 2018 Jan;45:176–80.

51. Kim JA, Kim HJ, Cho JM, Oh SH, Lee BH, Kim GH, et al. Diagnostic Value of Ceruloplasmin in the Diagnosis of Pediatric Wilson’s Disease. Pediatr Gastroenterol Hepatol Nutr. 2015 Sept;18(3):187–92.

52. Wang Y, Xu XL. Bacterial peptidoglycan-derived molecules activate Candida albicans hyphal growth. Commun Integr Biol. 2008;1(2):137–9.

53. Peters BM, Ovchinnikova ES, Krom BP, Schlecht LM, Zhou H, Hoyer LL, et al. Staphylococcus aureus adherence to Candida albicans hyphae is mediated by the hyphal adhesin Als3p. Microbiol Read Engl. 2012 Dec;158(Pt 12):2975–86.

54. Allison DL, Scheres N, Willems HME, Bode CS, Krom BP, Shirtliff ME. The Host Immune System Facilitates Disseminated Staphylococcus aureus Disease Due to Phagocytic Attraction to Candida albicans during Coinfection: a Case of Bait and Switch. Infect Immun. 2019 Nov;87(11):e00137–19.

55. Lu Y, Su C, Solis NV, Filler SG, Liu H. Synergistic regulation of hyphal elongation by hypoxia, CO(2), and nutrient conditions controls the virulence of Candida albicans. Cell Host Microbe. 2013 Nov 13;14(5):499–509.

56. Veiga H, Jousselin A, Schäper S, Saraiva BM, Marques LB, Reed P, et al. Cell division protein FtsK coordinates bacterial chromosome segregation and daughter cell separation in Staphylococcus aureus. EMBO J. 2023 June 1;42(11):e112140.

57. Gutgsell NS, Deutscher MP, Ofengand J. The pseudouridine synthase RluD is required for normal ribosome assembly and function in Escherichia coli. RNA N Y N. 2005 July;11(7):1141–52.

58. Dey S, Bishayi B. Riboflavin along with antibiotics balances reactive oxygen species and inflammatory cytokines and controls Staphylococcus aureus infection by boosting murine macrophage function and regulates inflammation. J Inflamm Lond Engl. 2016;13:36.

59. Ghssein G, Brutesco C, Ouerdane L, Fojcik C, Izaute A, Wang S, et al. Biosynthesis of a broad-spectrum nicotianamine-like metallophore in Staphylococcus aureus. Science. 2016 May 27;352(6289):1105–9.

60. Grim KP, San Francisco B, Radin JN, Brazel EB, Kelliher JL, Párraga Solórzano PK, et al. The Metallophore Staphylopine Enables Staphylococcus aureus To Compete with the Host for Zinc and Overcome Nutritional Immunity. mBio. 2017 Oct 31;8(5):e01281–17.

61. Duthie ES, Lorenz LL. Staphylococcal coagulase; mode of action and antigenicity. J Gen Microbiol. 1952 Feb;6(1–2):95–107.

62. Heurlier K, Dénervaud V, Pessi G, Reimmann C, Haas D. Negative control of quorum sensing by RpoN (sigma54) in Pseudomonas aeruginosa PAO1. J Bacteriol. 2003 Apr;185(7):2227–35.

63. Lew-Smith J, Binkley J, Sherlock G. The Candida Genome Database: annotation and visualization updates. Genetics. 2025 Mar 17;229(3):iyaf001.

64. Charpentier E, Anton AI, Barry P, Alfonso B, Fang Y, Novick RP. Novel cassette-based shuttle vector system for gram-positive bacteria. Appl Environ Microbiol. 2004 Oct;70(10):6076–85.

65. Stapels DAC, Ramyar KX, Bischoff M, von Köckritz-Blickwede M, Milder FJ, Ruyken M, et al. Staphylococcus aureus secretes a unique class of neutrophil serine protease inhibitors. Proc Natl Acad Sci U S A. 2014 Sept 9;111(36):13187–92.

66. Dalton T, Dowd SE, Wolcott RD, Sun Y, Watters C, Griswold JA, et al. An in vivo polymicrobial biofilm wound infection model to study interspecies interactions. PloS One. 2011;6(11):e27317.

67. Douglas EJA, Palk N, Brignoli T, Altwiley D, Boura M, Laabei M, et al. Extensive remodelling of the cell wall during the development of Staphylococcus aureus bacteraemia. eLife. 2023 July 4;12:RP87026.

68. Bae T, Schneewind O. Allelic replacement in Staphylococcus aureus with inducible counter-selection. Plasmid. 2006 Jan;55(1):58–63.

69. Monk IR, Tree JJ, Howden BP, Stinear TP, Foster TJ. Complete Bypass of Restriction Systems for Major Staphylococcus aureus Lineages. mBio. 2015 May 26;6(3):e00308–00315.

70. Laabei M, Uhlemann AC, Lowy FD, Austin ED, Yokoyama M, Ouadi K, et al. Evolutionary Trade-Offs Underlie the Multi-faceted Virulence of Staphylococcus aureus. PLoS Biol. 2015;13(9):e1002229.

71. Painter KL, Hall A, Ha KP, Edwards AM. The Electron Transport Chain Sensitizes Staphylococcus aureus and Enterococcus faecalis to the Oxidative Burst. Infect Immun. 2017 Dec;85(12):e00659–17.

