## Supplementary material for "Copper as a central element critical to interkingdom interactions within a polymicrobial wound environment"

**SUPPLMENTARY FIGURES AND TABLES**

**Table S1 –** CC22 GWAS hits for alleles associated with altered ability to persist in co-culture with *P. aeruginosa* and *C. albicans* using a wound-like model.

**Figure S1 –** Validation of CC22 GWAS hits using NTML transposon mutants to screen for impact on CFU recovery in mono (A) and polymicrobial (B) WLM models.

**Table S2–** *C. albicans* and *P. aeruginosa* CFU recovery following time-course assays in WLM polymicrobial community when complemented with GWAS hits (associated with Figure 2C-H).

**Table S3 –** Associated metadata for clinical *S. aureus* CC22 bloodstream isolates containing *copA* 1G>A mutation.

**Figure S2 –** JE2 and Newman copper exporter mutants time-course cultured with varying community structures

**Figure S3 –** Copper accumulation in JE2 copper exporter mutants when cultured in WLM and WLM-Cp**.**

**Figure S4 –** *Candida* driven copper accumulation sensitises *S. aureus* to ROS/RNS stress.

**Figure S5 –** Loss of copper exporter does not impact *S. aureus* sensitivity to pyocyanin, KCN and HQNO when cultured as a mono-species biofilm in WLM or WLM-CP.

**Figure S6 –** Loss of copper exporter does not impact competitiveness of *S. aureus* as a single-species culture in WLM or WLM-CP.

**Table S1 – CC22 GWAS hits for alleles associated with altered ability to persist in co-culture with *P. aeruginosa* and *C. albicans* using a wound-like model.**

| **SNP position** | **CC22 locus tag** | **Description** | **Log10P** | **NTML** |
| --- | --- | --- | --- | --- |
| 1485787 | SAEMRSA15_13160 | *dinG* | 3.922519 | NE346 |
| 2516262 | SAEMRSA15_23320 | putative glycerate kinase | 2.87224 | NE465 |
| 1240413 | SAEMRSA15_11090 | *ftsK* | 2.809844 | NE348 |
| 1996419 | SAEMRSA15_18100 | putative lipoprotein | 2.76335 | NE1087 |
| 830176 | SAEMRSA15_07180 | *nuc* | 2.62147 | NE1241 |
| 2515404 | SAEMRSA15_23310 | putative membrane protein | 2.507363 | NE602 |
| 889042 | SAEMRSA15_07880 | *gudB* | 2.438912 | NE1518 |
| 1130760 | SAEMRSA15_10160 | *murD* | 2.437838 | N/A |
| 448911 | SAEMRSA15_03860 | *metB* | 2.434957 | NE602 |
| 2758770 | SAEMRSA15_25490 | putative exported protein | 2.337558 | NE1353 |
| 1182957 | SAEMRSA15_10630 | *fabD* | 2.312296 | N/A |
| 1262036 | SAEMRSA15_11300 | *mutS* | 2.297622 | NE974 |
| 2554304 | SAEMRSA15_23640 | *cntB* | 2.297622 | NE1209 |
| 1566333 | Intergenic_Region_336 | Intergenic | 2.267633 | N/A |
| 2656043 | SAEMRSA15_24600 | *copA* | 2.267633 | NE561 |
| 977070 | Intergenic_Region_227 | Intergenic | 2.267633 | N/A |
| 661898 | SAEMRSA15_05660 | *tagX* | 2.267633 | NE1055 |
| 2441578 | SAEMRSA15_22660 | *mqo1* | 2.267633 | NE1033 |
| 844980 | SAEMRSA15_07420 | putative lipoprotein | 2.267633 | NE1589 |
| 1747240 | SAEMRSA15_15930 | *thrS* | 2.267633 | N/A |
| 1702225 | SAEMRSA15_15480 | *hisS* | 2.162897 | N/A |
| 590633 | SAEMRSA15_04910 | putative glycosyltransferase | 2.099388 | NE381 |
| 2577780 | SAEMRSA15_23880 | putative helicase | 2.095475 | NE513 |
| 893074 | SAEMRSA15_07910 | *argG* | 2.090088 | NE505 |
| 1697502 | SAEMRSA15_15450 | putative ATPase | 2.069294 | NE341 |
| 506893 | Intergenic_Region_105 | Intergenic | 1.944719 | N/A |
| 447344 | SAEMRSA15_03840 | sodium:neurotransmitter symporter family protein | 1.715478 | NE231 |
| 2560821 | SAEMRSA15_23690 | hypothetical protein | 1.697099 | NE633 |
| 1856658 | SAEMRSA15_16750 | *ribBA* | 1.691634 | NE573 |
| 2538962 | SAEMRSA15_23500 | putative amino acid permease | 1.685415 | NE1131 |
| 2325657 | SAEMRSA15_21480 | *rplD* | 1.68173 | N/A |
| 2746102 | SAEMRSA15_25410 | *argR3* | 1.654027 | NE367 |
| 566273 | SAEMRSA15_04760 | putative peptidase | 1.581785 | NE1455 |
| 149065 | SAEMRSA15_01210 | *capG5* | 1.573339 | NE580 |
| 2617270 | SAEMRSA15_24210 | nitroreductase family protein | 1.527706 | NE441 |
| 1145714 | SAEMRSA15_10300 | *rluD* | 1.527706 | NE793 |
| 2802186 | SAEMRSA15_25850 | *hisG* | 1.519118 | NE103 |
| 2385067 | SAEMRSA15_22080 | *lcpC* | 1.482041 | NE415 |
| 1479055 | SAEMRSA15_13110 | *pbp2* | 1.465598 | N/A |
| 2292604 | Intergenic_Region_513 | Intergenic | 1.433578 | N/A |
| 342920 | SAEMRSA15_02780 | NADH:flavin oxidoreductase / NADH oxidase family protein | 1.330221 | NE1199 |
| 2312449 | SAEMRSA15_21210 | *cbiO* | 1.327985 | NE1148 |

**Figure S1 – Validation of CC22 GWAS hits using NTML transposon mutants to screen for impact on CFU recovery in mono (A) and polymicrobial (B) WLM models.** No significant difference detected in GWAS hits when cultured as *S. aureus* biofilm in WLM at the 3-day timepoint (A). When cultured as tri-species biofilms ftsK, copA, ribBA, rluD, camS and cntB were negatively impacted whilst transposon mutant strains NE1753 and NE1782 demonstrated increased persistence (B) Data collated from three independent experiments with points representing mean of each. At each timepoint a minimum of three biofilms were processed and CFU’s enumerated. Bars represent the mean value, error bars the standard deviation. Significance determined as **<0.01, ***<0.001, ****<0.0001 calculated using a one-way ANOVA with Dunnett’s multiple comparisons test.

A

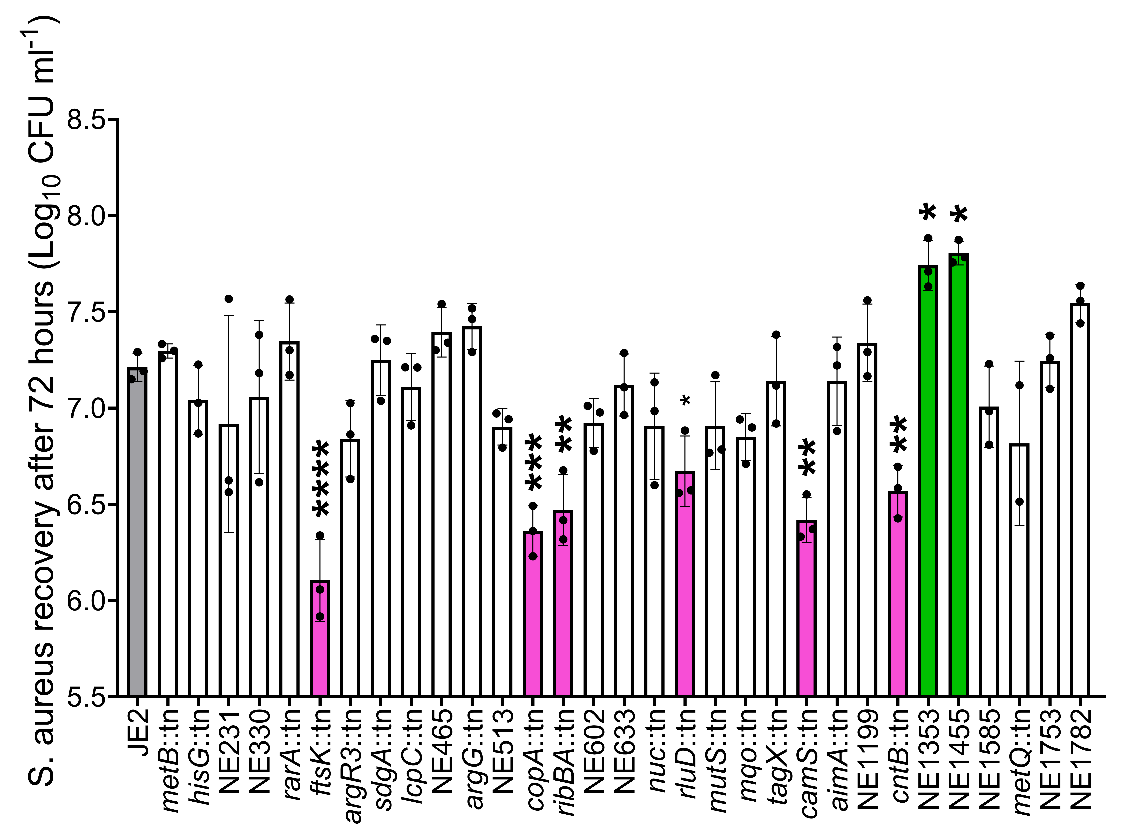

B

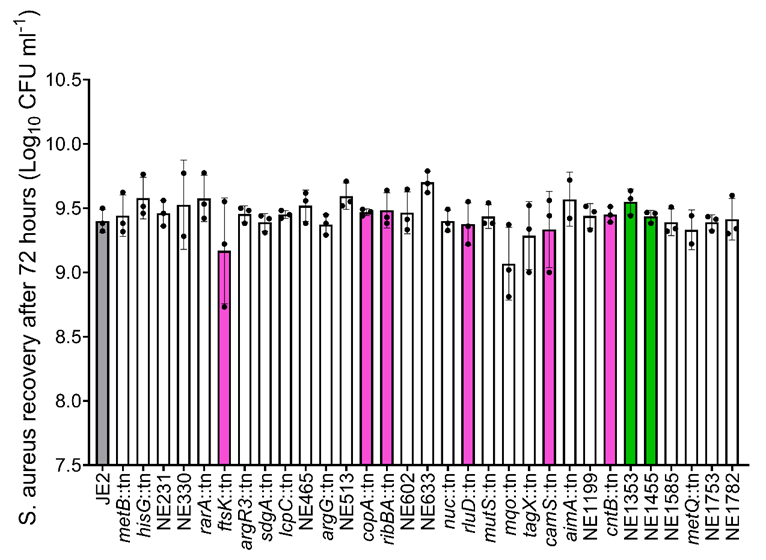

**Table S2– *C. albicans* and *P. aeruginosa* CFU recovery following time-course assays in WLM polymicrobial community when complemented with GWAS hits (associated with Figure 2C-H).** Average represents three individual experiments with at least three biofilms sacrificed for enumeration at the relevant timepoints. Significant findings highlighted in bold (* <0.05)

| ***S. aureus* strains** | ***C. albicans* CFU (log_10_ ml^-1^)** | | | ***P. aeruginosa* CFU (log_10_ ml^-1^)** | | |
| --- | --- | --- | --- | --- | --- | --- |
|  | **Average** | **St.dev** | **N** | **Average** | **St.dev** | **N** |
| JE2 wild type | 5.18 | 0.22 | 3 | 9.37 | 0.35 | 3 |
| *copA*::tn | 5.32 | 0.21 | 3 | 9.27 | 0.23 | 3 |
| *copA*::tn pCN34 | 4.93 | 0.22 | 3 | 9.05 | 0.44 | 3 |
| *copA*::tn p*copA* | 5.03 | 0.23 | 3 | 9.21 | 0.21 | 3 |
| ***ftsK*::tn** | **5.73** | **0.15** | **3** | **9.87** | **0.152753** | **3** |
| ***ftsK*::tn pCN34** | **5.64** | **0.32** | **3** | **9.93** | **0.42** | **3** |
| *ftsK*::tn p*ftsK* | 4.91 | 0.44 | 3 | 9.28 | 0.38 | 3 |
| *ribBA*::tn | 5.22 | 0.28 | 3 | 9.18 | 0.42 | 3 |
| *ribBA*::tn pCN34 | 4.95 | 0.11 | 3 | 9.42 | 0.61 | 2 |
| *ribBA*::tn p*ribBA* | 5.33 | 0.32 | 3 | 9.28 | 0.28 | 2 |
| *cntB*::tn | 5.41 | 0.28 | 3 | 9.57 | 0.41 | 3 |
| *cntB*::tn pCN34 | 5.28 | 0.40 | 3 | 9.62 | 0.33 | 3 |
| *cntB*::tn p*cntB* | 5.18 | 0.15 | 3 | 9.21 | 0.18 | 3 |
| *rluD*::ten | 5.21 | 0.43 | 3 | 9.50 | 0.19 | 3 |
| *rluD*::tn pCN34 | 5.05 | 0.41 | 3 | 9.32 | 0.47 | 3 |
| *rluD*::tn p*rluD* | 4.92 | 0.21 | 3 | 9.11 | 0.22 | 3 |
| 2579::tn | 4.91 | 0.48 | 3 | 9.15 | 0.28 | 3 |
| 2579::tn pCN34 | 4.82 | 0.37 | 2 | 9.02 | 0.39 | 3 |
| 2579::tn p2579 | 5.21 | 0.22 | 3 | 9.32 | 0.3 | 3 |

**Table S3 – Associated metadata for clinical *S. aureus* CC22 bloodstream isolates containing *copA* 1G>A mutation.** For CC22 strains containing *copA* 1G>A mutation, 8 of 10 initially entered the patient as a soft tissue infection. This may represent a niche specific adaptation which improves survival for S. aureus in wound environments. The ability to successfully compete in this niche enables S. aureus to persist for longer, potentially increasing chances of progression to bloodstream infection.

| **Strain ID** | **Date of isolation** | **Collection** | **Entry point** | **Classification** | **Outcome (90 days)** | **WLM CFU recovery (Log_10_)** |
| --- | --- | --- | --- | --- | --- | --- |
| ASARM76 | 9/28/2006 | MRSA | SoftTissueInfection | HospitalAcquired | Death | 7.45 |
| ASARM101 | 1/30/2007 | MRSA | SoftTissueInfection | HealthCareAssociated | Death | 5.89 |
| ASARM138 | 9/15/2007 | MRSA | Line | HospitalAcquired | Death | 7.56 |
| ASARM165 | 8/11/2008 | MRSA | SoftTissueInfection | HealthCareAssociated | Alive | 6.94 |
| ASARMLT1 | 8/14/2008 | MRSA | SoftTissueInfection | HealthCareAssociated | Alive | 7.74 |
| ASARMLT2 | 10/5/2008 | MRSA | SoftTissueInfection | HealthCareAssociated | Alive | 7.68 |
| ASARMLT3 | 11/3/2008 | MRSA | SoftTissueInfection | HealthCareAssociated | Alive | 7.80 |
| ASARM170 | 9/29/2008 | MRSA | SoftTissueInfection | HealthCareAssociated | Alive | 7.39 |
| ASARM171 | 10/20/2008 | MRSA | SoftTissueInfection | HealthCareAssociated | Alive | 7.49 |
| ASASM132 | 11/15/2007 | MSSA | DeepTissueAbcess | HospitalAcquired | Alive | 7.48 |

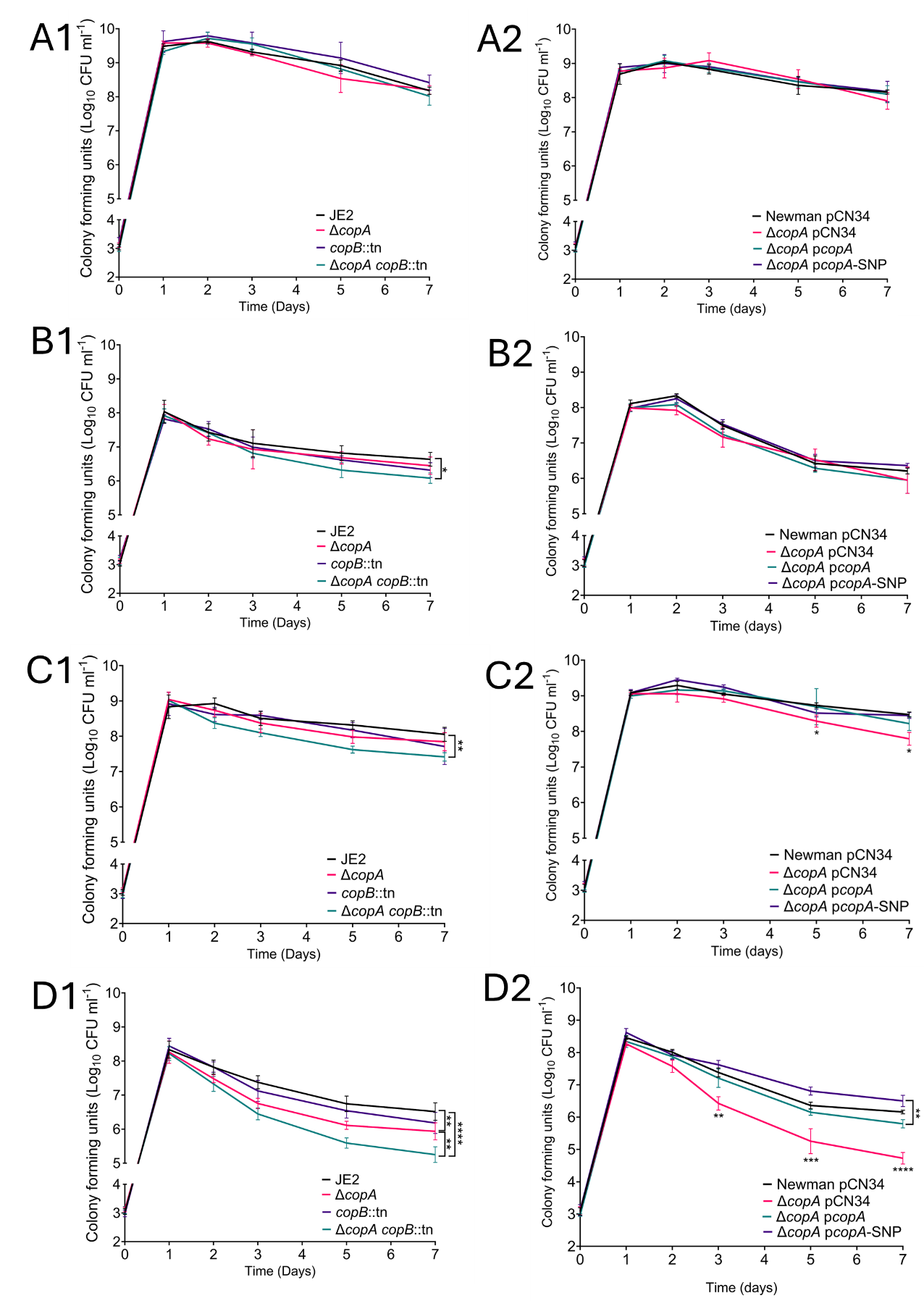
**Figure S2 – JE2 and Newman copper exporter mutants time-course cultured with varying community structures.** Panels with ‘1’ denote JE2 based strains, panels with ‘2’ denote Newman based strains. A) *S. aureus* mono-species biofilms in WLM. B) *S. aureus*/ *P. aeruginosa* dual-species biofilms in WLM. C) S. aureus/C. albicans dual-species biofilms in WLM. D) *S. aureus*/*P. aeruginosa*/ *C. albicans* tri-species biofilms in WLM. CFU’s enumerated following incubation at 37^O^C for defined period with at least 3 biofilms sacrificed at each time point. Data collated from three independent experiments with points representing mean of each. Bars represent the mean value, error bars the standard deviation. Significance determined as *<0.05, **<0.01, ***<0.001, ****<0.0001 calculated using a two-way ANOVA with Tukey’s multiple comparisons test.

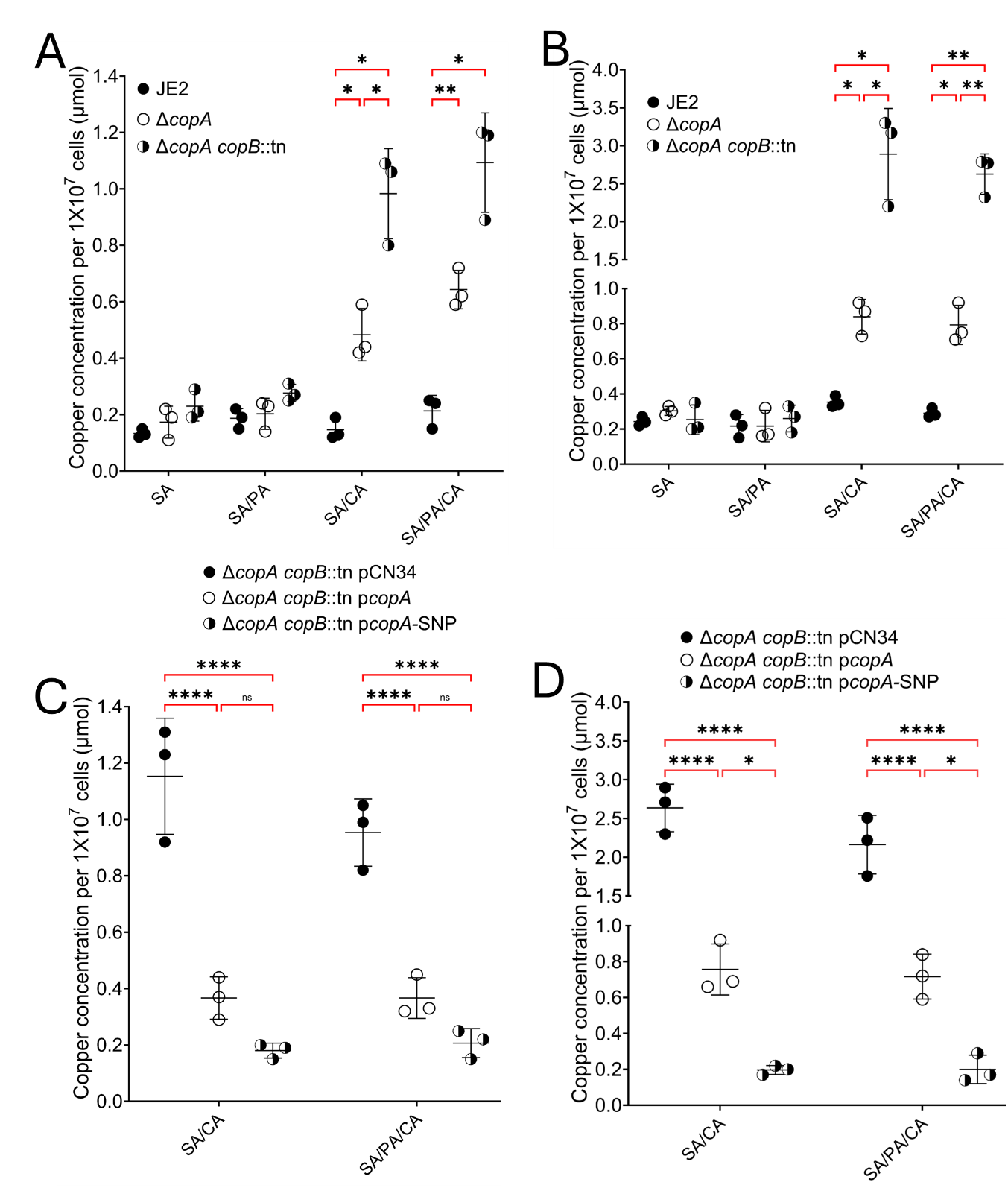
**Figure S3 – Copper accumulation in JE2 copper exporter mutants when cultured in WLM and WLM-Cp.** Intracellular contents of JE2 derived *S. aureus* strains liberated through lysostaphin digestion after biofilm disruption. WLM biofilms established as mono, dual and tri-species cultures in WLM only (A) and WLM+20µM human ceruloplasmin (B). Impact of complementation with the wild type *copA* allele and SNP in WLM (C) or WLM+20µM human ceruloplasmin (D). Copper only accumulates in JE2 ∆*copA* when cultured in the presence of *C. albicans,* with even higher levels of copper accumulating in ∆*copA copB*::tn (A). Addition of exogenous human ceruloplasmin increases the level of copper accumulation in copper exporter mutants, suggesting *C. albicans* is liberating copper from ceruloplasmin (B). Restoration of the wild type *copA* allele reduced copper accumulation in JE2 ∆*copA copB::*tn (C), with complementation of *copA* G>A1 further reducing copper accumulation (D). Data collated from three independent experiments with points representing mean of each. Lines represent the mean value, error bars the standard deviation. Significance determined as *<0.05, **<0.01, ***<0.001, ****<0.0001 calculated using Sidak’s multiple comparisons test.

**Figure S4 – *Candida* driven copper accumulation sensitises *S. aureus* to ROS/RNS stress.** Hydrogen peroxide killing assay of strains cultured in WLM or WLM-CP (A). Comparison within WLM or WLM-CP indicated in black. Comparison between WLM and WLM-CP indicated in red. Deletion of *copA* sensitises *S. aureus* to H_2_O_2_ due to accumulation of copper. In WLM ceruloplasmin is derived from plasma, whilst increasing ceruloplasmin through exogenous addition increases the sensitivity of ∆*copA* to H_2_O_2_ mediated killing. Complementation restores killing phenotype with restoration of *copA* 1G>A providing enhanced protection to H_2_O_2_ through minimising intracellular copper. Thiourea scavenging of ROS protects JE2 ∆*copA* from toxic effects of copper accumulation (B). Pre-incubation of cells with thiourea protects *S. aureus* from H_2_O_2_ toxicity. Enhanced sensitivity to H_2_O_2_ through cytoplasmic copper accumulation lost in JE2 ∆*copA.* Strains cultured as a tri-species biofilm in WLM for all experiments*.* Human Ceruloplasmin added to media at 20 µM where indicated. CFU’s enumerated following incubation of 1x10^6^ cells in 20 mM H_2_O_2_ for 3 H. Data collated from three independent experiments, symbols represent the mean value, error bars the standard deviation. Significance determined as *<0.05, **<0.01, ***<0.001, ****<0.0001 calculated using a two-way Anova with Tukey’s post-hoc test (A) and Three-way Anova with Sidak’s multiple comparisons test (B).

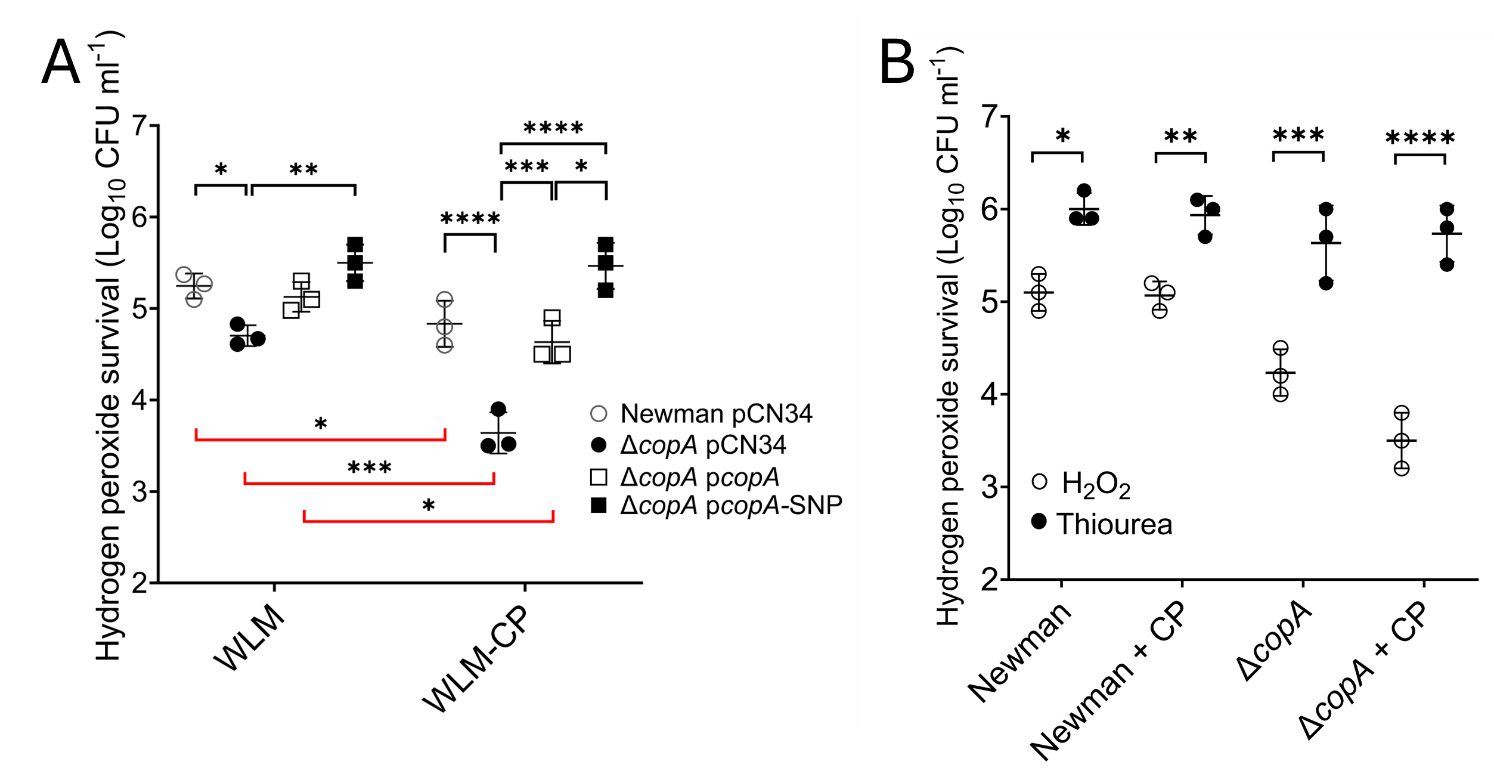

**Figure S5 – Loss of copper exporter does not impact *S. aureus* sensitivity to pyocyanin, KCN and HQNO when cultured as a mono-species biofilm in WLM or WLM-CP.** Mono-species biofilms cultured in WLM or WLM-CP for 3 days. Cells normalised to 1X10^6^ and exposed to 0.2 mM Pyocyanin, 1 mM KCN and 0.2 mM HQNO for 6 H, followed by plating for CFU. When cultured as a mono-species biofilms, activity of pyocyanin, KCN and HQNO is not enhanced by the presence of ceruloplasmin. In the absence of *C. albicans* mediated release of copper from ceruloplasmin, these *P. aeruginosa* derived antimicrobials maintain the same level of effectiveness irrespective of strain background or supplementation of ceruloplasmin. Data collated from three independent experiments, symbols represent the mean value, error bars the standard deviation. CFU enumeration found to not be significant using a three-way ANOVA with Sidaks’ post hoc analysis.

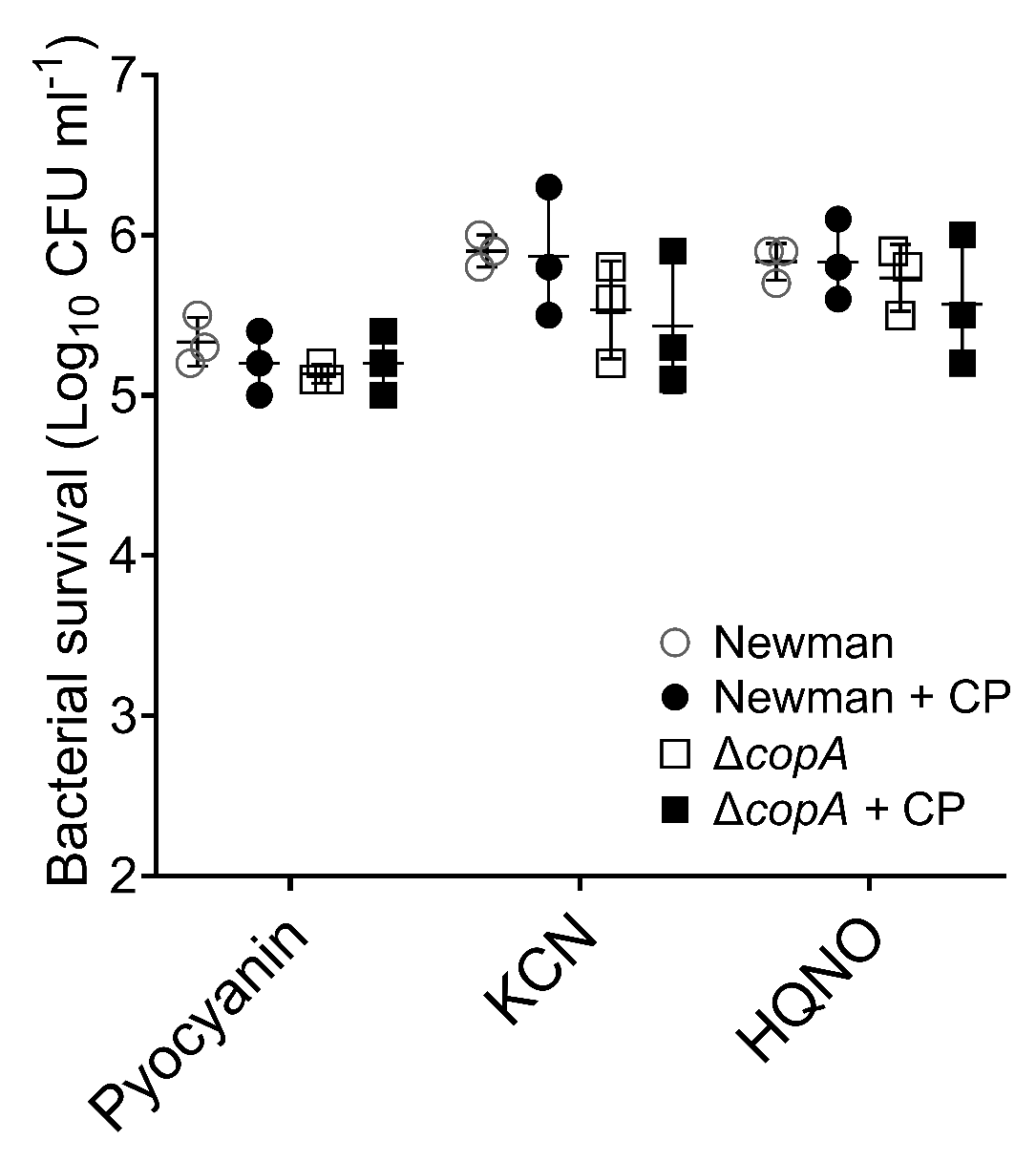

**Figure S6 – Loss of copper exporter does not impact competitiveness of *S. aureus* as a single-species culture in WLM or WLM-CP.** S. aureus only WLM and WLM-Cp biofilms set up and incubated for 3 days. In the absence of *C. albicans*, copper is not released from ceruloplasmin and does not accumulate in copper exporter mutants. No competitive defect is detected when compared to a rifampicin resistant marker strain for a *copA* mutant, with no benefit for strains expressing *copA* 1G>A. Data collated from three independent experiments with points representing mean of each. Lines represent the mean value and error bars the standard deviation. Competition index found to not be significant using a two-way ANOVA with Tukey’s post hoc analysis.

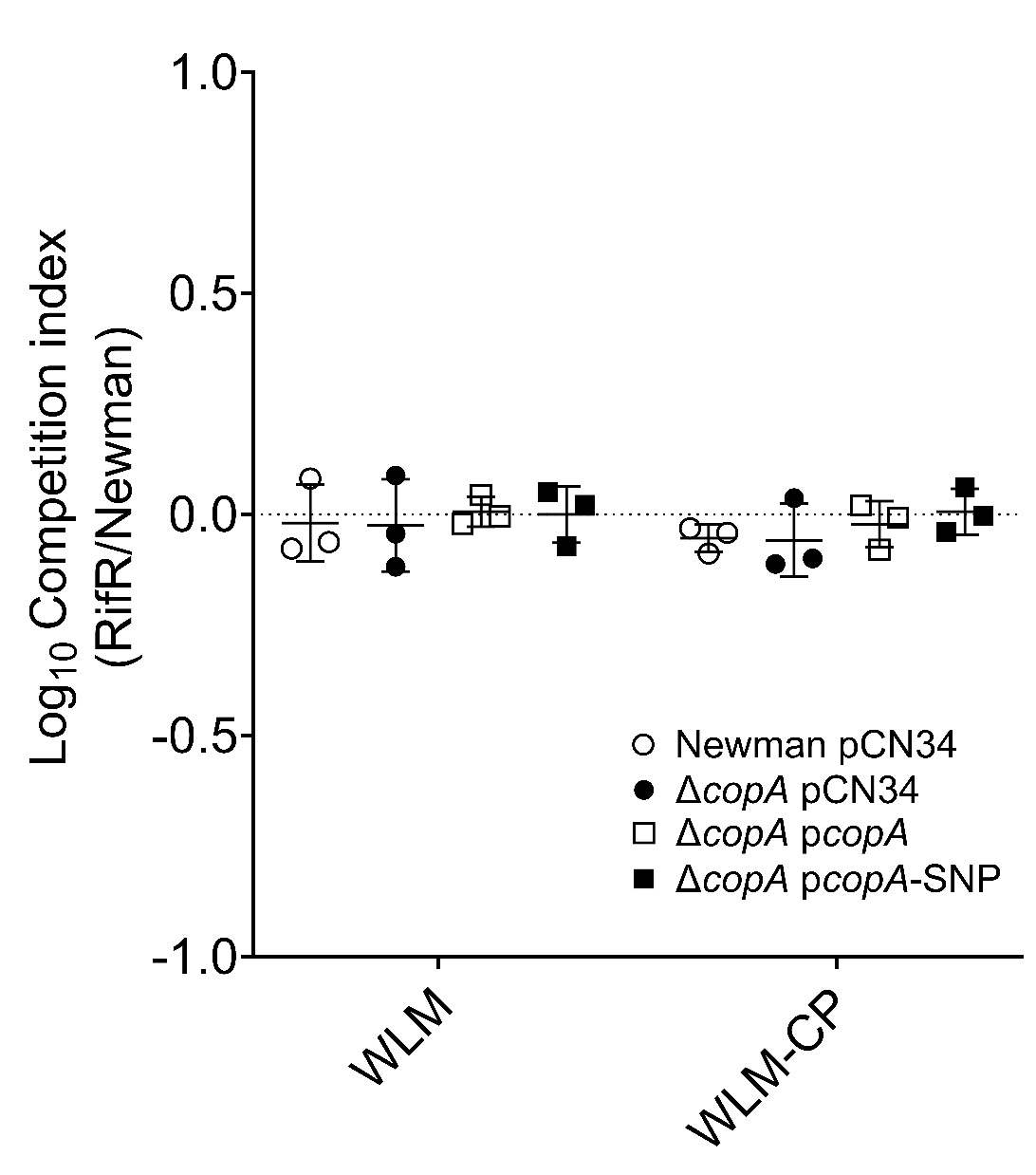
